# Key features of the genetic architecture and evolution of host-microbe interactions revealed by high-resolution genetic mapping of the mucosa-associated gut microbiome in hybrid mice

**DOI:** 10.1101/2021.09.28.462095

**Authors:** Shauni Doms, Hanna Fokt, Malte Christoph Rühlemann, Cecilia J. Chung, Axel Künstner, Saleh Ibrahim, Andre Franke, Leslie M. Turner, John F. Baines

## Abstract

Determining the forces that shape diversity in host-associated bacterial communities is critical to understanding the evolution and maintenance of metaorganisms. To gain deeper understanding of the role of host genetics in shaping gut microbial traits, we employed a powerful genetic mapping approach using inbred lines derived from the hybrid zone of two incipient house mouse species. Further, we uniquely performed our analysis on microbial traits measured at the gut mucosal interface, which is in more direct contact with host cells and the immune system. A high number of mucosa-associated bacterial taxa have significant heritability estimates; heritabilities are greater for 16S rRNA transcript-compared to gene copy-based traits, and interestingly, are positively correlated with cospeciation rate estimates. Genomewide association mapping identifies 443 loci influencing 123 taxa, with narrow genomic intervals pinpointing promising candidate genes and pathways. Importantly, we identified an enrichment of candidate genes associated with several human diseases, including inflammatory bowel disease, and functional categories including innate immunity and G-protein-coupled receptors. These results highlight key features of the genetic architecture of mammalian host-microbe interactions and how they diverge as new species form.

## Introduction

The recent widespread recognition of the gut microbiome’s importance to host health and fitness represents a critical advancement of biomedicine. Host phenotypes affected by the gut microbiome are documented in humans (Ley et al., 2006; Turnbaugh et al., 2009; Lynch and Pedersen, 2016), laboratory animals (Backhed et al., 2004; Turnbaugh et al., 2008; Rolig et al., 2015; Rosshart et al., 2017; Gould et al., 2018), and wild populations (Suzuki, 2017; Roth et al., 2019; Suzuki et al., 2020; Hua et al., 2020), and include critical traits such as aiding digestion and energy uptake (Rowland et al., 2018), and the development and regulation of the immune system (Davenport, 2020).

Despite the importance of gut microbiome, community composition varies significantly among host species, populations, and individuals (Benson et al., 2010; Yatsunenko et al., 2012; Brooks et al., 2016; Rehman et al., 2016; Amato et al., 2019). While a portion of this variation is expected to be selectively neutral, alterations of the gut microbiome are on the one hand linked to numerous human diseases (Carding et al., 2015; Lynch and Pedersen, 2016) such as diabetes (Qin et al., 2012), inflammatory bowel disease (IBD) (Ott et al., 2004; Gevers et al., 2014) and mental disorders (Clapp et al., 2017). On the other hand, there is evidence that the gut microbiome can play an important role in adaptation on both recent- (Hehemann et al., 2010; Suzuki and Ley, 2020) and ancient evolutionary timescales (). Collectively, these phenomena suggest that it would be evolutionarily advantageous for hosts to influence their microbiome.

An intriguing observation made in comparative microbiome research in the last decade is that the pattern of diversification among gut microbiomes appears to mirror host phylogeny (Ochman et al., 2010). This phenomenon, coined “phylosymbiosis” (Brucker and Bordenstein, 2012a; Brucker and Bordenstein, 2012b; Lim and Bordenstein, 2020), is documented in a number of diverse host taxa (Brooks et al., 2016) and also extends to the level of the phageome (Gogarten et al., 2021). Several non-mutually exclusive hypotheses are proposed to explain phylosymbiosis (Moran and Sloan, 2015). However, it is likely that vertical inheritance is important for at least a subset of taxa, as signatures of co-speciation/-diversification are present among numerous mammalian associated gut microbes (Moeller et al., 2016; Groussin et al., 2017; Moeller et al., 2019), which could also set the stage for potential coevolutionary processes. Importantly, experiments involving interspecific fecal microbiota transplants indeed provide evidence of host adaptation to their conspecific microbial communities (Brooks et al., 2016; Moeller et al., 2019). Further, cospeciating taxa were observed to be significantly enriched among the bacterial species depleted in early onset IBD, an immune-related disorder, suggesting a greater evolved dependency on such taxa (Papa et al., 2012; Groussin et al., 2017). However, the nature of genetic changes involving host-microbe interactions that take place as new host species diverge remains under-explored.

House mice are an excellent model system for evolutionary microbiome research, as studies of both natural populations and laboratory experiments are possible (Suzuki, 2017; Suzuki et al., 2019). In particular, the house mouse species complex is comprised of subspecies that hybridize in nature, enabling the potential early stages of codiversification to be studied. We previously analyzed the gut microbiome across the central European hybrid zone of *Mus musculus musculus* and *M. m. domesticus* (Wang et al., 2015), which share a common ancestor ~ 0.5 million years ago (Geraldes et al., 2008). Importantly, transgressive phenotypes (i.e. exceeding or falling short of parental values) among gut microbial traits as well as increased intestinal histopathology scores were common in hybrids, suggesting that the genetic basis of host control over microbes has diverged (Wang et al., 2015). The same study performed an F_2_ cross between wild-derived inbred strains of *M. m. domesticus* and *M. m. musculus* and identified 14 quantitative trait loci (QTL) influencing 29 microbial traits. However, like classical laboratory mice, these strains had a history of rederivation and reconstitution of their gut microbiome, thus leading to deviations from the native microbial populations found in nature (Rosshart et al., 2017; Org and Lusis, 2018), and the genomic intervals were too large to identify individual genes.

In this study, we employed a powerful genetic mapping approach using inbred lines directly derived from the *M. m. musculus - M. m. domesticus* hybrid zone, and further focus on the mucosa-associated microbiota due to its more direct interaction with host cells (Fukata and Arditi, 2013; Chu and Mazmanian, 2013), distinct functions compared to the luminal microbiota (Wang et al., 2010; Vaga et al., 2020), and greater dependence on host genetics (Spor et al., 2011; Linnenbrink et al., 2013). Previous mapping studies using hybrids raised in a laboratory environment showed that high mapping resolution is possible due to the hundreds of generations of natural admixture between parental genomes in the hybrid zone (Turner and Harr, 2014; Pallares et al., 2014; Škrabar et al., 2018). Accordingly, we here identify 443 loci contributing to variation in 123 taxa, whose narrow genomic intervals (median <2Mb) enable many individual candidate genes and pathways to be pinpointed. We identify a high proportion of bacterial taxa with significant heritability estimates, and find that bacterial phenotyping based on 16S rRNA transcript compared to gene copy-based profiling yields an even higher proportion. Further, these heritability estimates also significantly positively correlate with cospeciation rate estimates, suggesting a more extensive host genetic architecture for cospeciating taxa. Finally, we identify numerous enriched functional pathways, whose role in host-microbe interactions may be particularly important as new species form.

## Results

### Microbial community composition

To obtain microbial traits for genetic mapping in the G2 mapping population, we sequenced the 16S rRNA gene from caecal mucosa samples of 320 hybrid male mice based on DNA and RNA (cDNA), which reflect bacterial cell number and activity, respectively. After applying quality filtering and subsampling 10,000 reads per sample, we identified a total of 4684 amplicon sequence variants (ASVs). For further analyses, we established a “core microbiome” (defined in Methods), such that analyses were limited to those taxa common and abundant enough to reveal potential genetic signal. The core microbiome is composed of four phyla, five classes, five orders, eleven families, 27 genera, and 90 ASVs for RNA, and four phyla, five classes, six orders, twelve families, 28 genera and 46 ASVs for DNA. A combined total of 98 unique ASVs belong to the core, of which 38 were shared between DNA and RNA (Suppl. Fig. 1). The most abundant genus in our core microbiome is *Helicobacter* (Suppl. Fig. 2), consistent with a previous study of the wild hybrid *M. m. musculus/M. m. domesticus* mucosa-associated microbiome (Wang et al., 2015).

### Correlation between host genetic relatedness and microbiome structure

To gain a broad sense of the contribution of genetic factors to the variability of microbial phenotypes in our mapping population, we compared the kinship matrix based on genotypes to an equivalent based on gut microbial composition, whereby ASV abundances were used as equivalents of gene dosage. We found a significant correlation between these matrices (*P* = .001; Suppl. Fig. 3), indicating a host genetic effect on the diversity of the gut microbiota.

### SNP-based heritability

Next, we used a SNP-based approach to estimate the proportion of variance explained (PVE) of the relative abundance of taxa, also called the narrow-sense heritability (*h*^2^) or SNP-based heritability. Out of the 153 total core taxa, we identified 46 taxa for DNA and 69 taxa for RNA with significant heritability estimates (*P_RLRT_* < .05), with estimates ranging between 29 and 91% (see Fig. 1A-B and Suppl. Table 1). An unclassified genus belonging to the phylum Bacteroidetes followed by ASV7 (genus *Paraprevotella), Paraprevotella* and Paraprevotellaceae showed the highest heritability among DNA-based traits (91.8%, 88.8%, 88.8%, and 87.1%, respectively; Fig. 1A), while ASV97 (genus *Oscillibacter*), followed by Prevotellaceae, *Paraprevotella* and ASV7 (*Paraprevotella*) had the highest heritability among RNA-based traits (86.6%, 85.7%, 85.7%, and 85.6%, resp.; Fig. 1B). The heritability estimates for DNA- and RNA-based measurements of the same taxa are significantly correlated (*P* = 5.013 x 10^-8^, R^2^=0.58, Suppl. Fig. 4), and neither measure appears to be systematically more heritable than another, i.e. some taxa display higher RNA-based heritability estimates and others higher DNA-based estimates.

**Figure 1:**
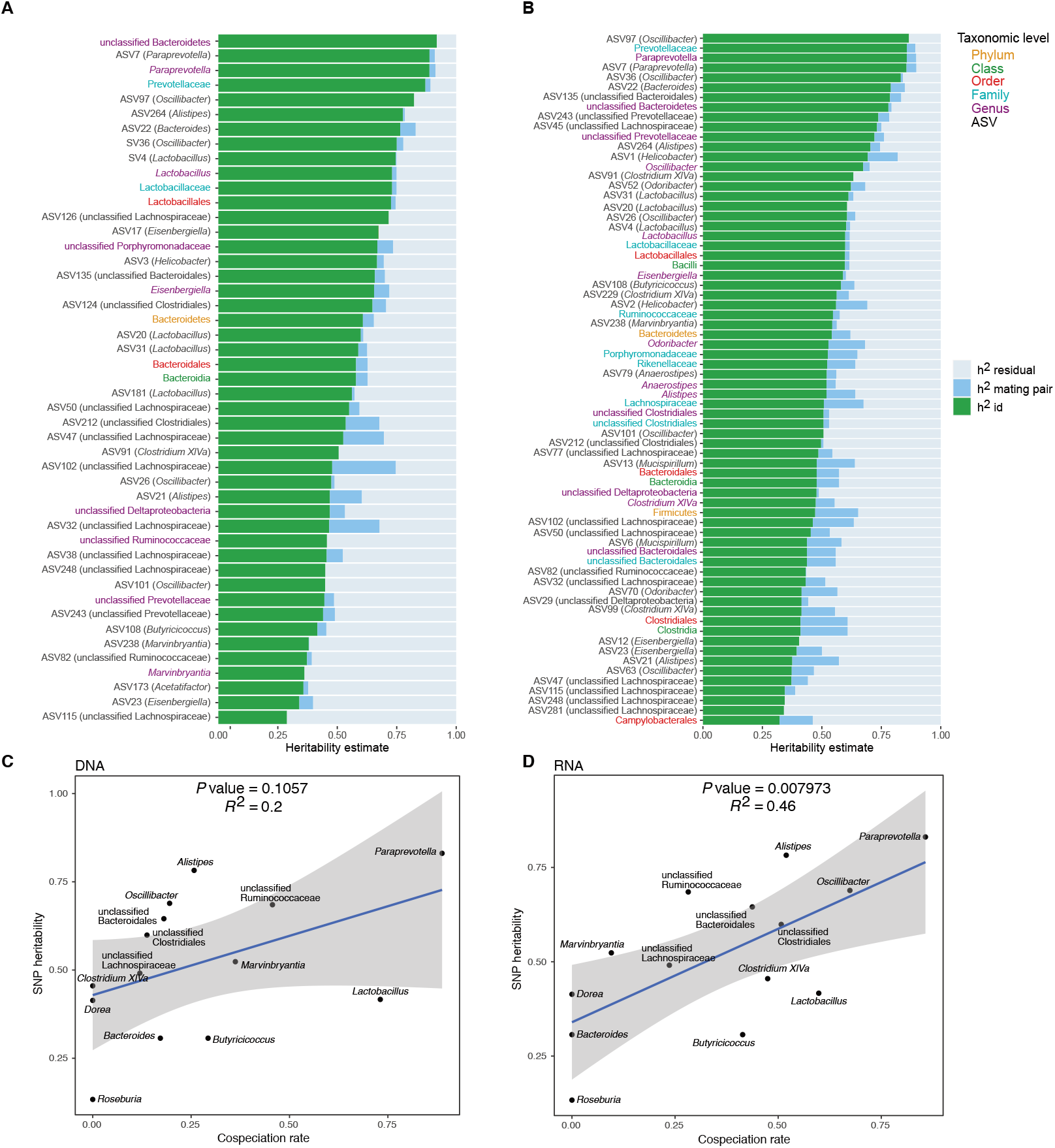
(A-B) Heritability estimates for the relative abundance of bacterial taxa. Proportion of variance explained for each taxon on DNA level (A), and RNA level (B) for all SNPs (GRM) in green, mating pair identifier in blue and residual variance in grey. Only significant heritability estimates are shown (*P* < .05). The text labels on the y-axis are colored according to taxonomic level: ASV in black, genus in purple, family in light blue, order in red, class in green, and phylum in yellow. (C-D) Relationship between the heritability estimates for the relative abundance of bacterial taxa and co-speciation rate for the same genus calculated by Groussin et al. (2017). DNA level (C), and RNA level (D). The blue line represents a linear regression fit to the data and the grey area the corresponding confidence interval.

### Heritability estimates are correlated with predicted co-speciation rates

In an important meta-analysis of the gut microbiome across diverse mammalian taxa, Groussin et al. (2017) estimated co-speciation rates of individual bacterial taxa by measuring the congruence of host and bacteria phylogenetic trees relative to the number of host-swap events. We reasoned that taxa with higher co-speciation rates might also demonstrate higher heritability, as these more intimate evolutionary relationships would provide a greater opportunity for genetic aspects to evolve. Intriguingly, we observe a significant positive correlation for RNA-based traits (*P*= .008, R^2^=.46, Fig. 1D) and a similar trend for DNA (*P*= 0.1; Fig. 1C). These results support the notion that cospeciating taxa evolved a greater dependency on host genes, and further suggest that bacterial activity may better reflect the underlying biological interactions.

### Genetic mapping of host loci determining microbiome composition

Next, we performed genome-wide association mapping of the relative abundances of core taxa, in addition to two alpha-diversity measures (Shannon and Chao1 indices), based on 32,625 SNPs. We included both additive and dominance terms in the model to enable the identification of under- and over-dominance (see Methods). While we found no significant associations for alpha diversity at either the DNA or RNA level (*P* > 1.53 × 10^-6^), a total of 1099 genome-wide significant associations were identified for individual taxa (*P* < 1.53 × 10^-6^, Suppl. Table 2), of which 443 achieved study-wide significance (*P* < 1.29 × 10^-8^). Apart from the X chromosome, all autosomal chromosomes contained study-wide significant associations (Fig. 2). Out of the 153 mapped taxa, 123 had at least one significant association (Table 1). For the remainder of our analyses, we focus on the results using the more stringent study-wide threshold, and combined significant SNPs within 10 Mb into significant regions (Suppl. Table 3). The median size of significant regions is 1.91 Mb, which harbor a median of 14 protein-coding genes. On average, we observe 10 significant mouse genomic regions per bacterial taxon.

**Figure 2:**
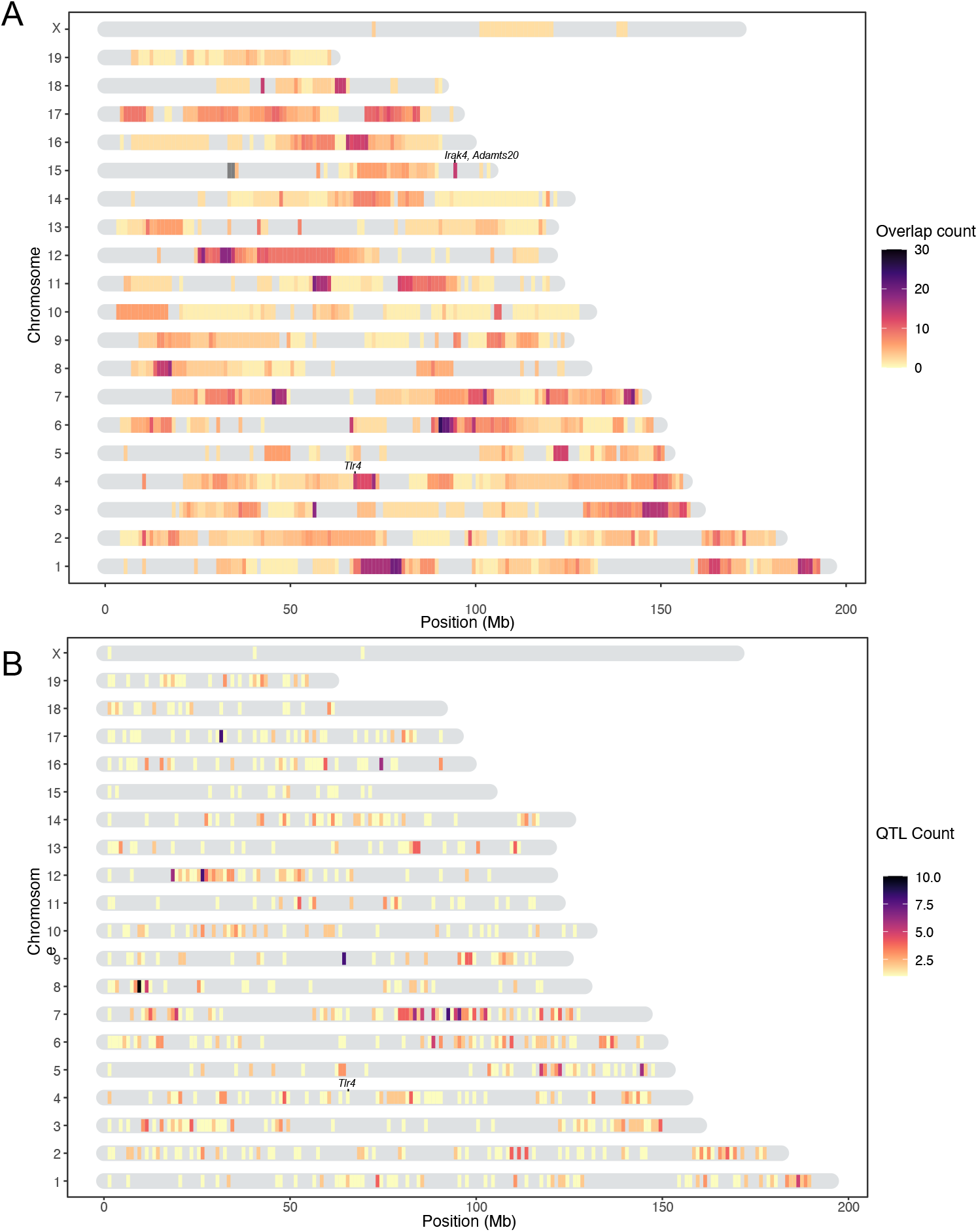
Heatmap of significant host loci from association mapping of bacterial abundances. Karotype plot showing the number of significant loci found using a studywide threshold, where (A) plots the significance intervals, and (B) the significant SNP markers on the chromosomes.

**Table. 1.** Overview of mapping statistics.

|  | DNA | RNA | Total |
| --- | --- | --- | --- |
| Mapped taxa | 101 | 142 | 153 |
| Taxa with significant loci | 67 | 96 | 123 |
| Median interval size (Mb) | 1.52 | 2.29 | 1.91 |
| Total significant loci | 478 | 791 | 1269 |
| Unique significant loci | 179 | 313 | 443 |
| Significant loci total $P$ | 91 | 167 | 233 |
| Significant loci additive $P$ | 155 | 260 | 377 |
| Significant loci dominance $P$ | 95 | 166 | 231 |
| Median significant loci per trait | 5 | 6 | 8 |
| Median unique significant loci per trait | 3 | 3 | 4 |
| Median unique significant SNPs per locus | 2 | 2.5 | 2 |
| Median number of genes per locus | 31 | 52 | 43 |
| Median protein coding genes per locus | 11 | 15 | 14 |

Of the significant loci with estimated interval sizes, we find 73 intervals (16.5%) that are smaller than one Mb (Suppl. Table 4). The smallest interval is only 231 bases and associated with the RNA-based abundance of an unclassified genus belonging to Deltaproteobacteria. It is situated in an intron of the C3 gene, a complement component playing a central role in the activation of the complement system, which modulates inflammation and contributes to antimicrobial activity (Ricklin et al., 2016).

The significant genomic regions and SNPs are displayed in Figure 2A and 2B, respectively. Individual SNPs were associated with up to 12 taxa, and significant intervals with up to 30 taxa. The SNPs with the lowest *P* values were associated with the genus *Dorea* and two ASVs belonging to *Dorea* (ASV184 and ASV293; Suppl. Fig. 5). At the RNA level this involves two loci: mm10-chr4: 67.07 Mb, where the peak SNP is 13 kb downstream of the closest gene *Tlr4* (UNC7414459, *P*=2.31 × 10^-69^, additive *P*= 4.48 × 10^-118^, dominance *P*= 1.37 × 10^-111^), and mm10-chr15: 94.4 Mb, where the peak SNP is found within the *Adamts20* gene (UNC26145702, *P*=4.51 × 10^-65^, additive *P*= 1.87 × 10^-113^, dominance *P*= 1.56 × 10^-105^; Fig. 2; Suppl. Fig. 5). Interestingly, the *Irak4* gene, whose protein product is rapidly recruited after TLR4 activation, is also located 181 kb upstream of *Adamts20*. The five taxa displaying the most associations were ASV19 (*Bacteroides*), *Dorea*, ASV36 (*Oscillibacter*), ASV35 (*Bacteroides*), and ASV98 (unclassified Lachnospiraceae) (Suppl. Fig. 6).

### Ancestry, dominance, and effect sizes

A total of 435 significant SNPs were ancestry informative between *M. m. musculus* and *M. m. domesticus (i.e.* represent fixed differences between subspecies). To gain further insight on the genetic architecture of microbial trait abundances, we estimated the degree of dominance at each significant locus using the *d/a* ratio (Falconer, 1996), where alleles with strictly recessive, additive, and dominant effects have *d/a* values of −1, 0, and 1, respectively. As half of the SNPs were not ancestry informative (Fig. 3A), it was not possible to consistently have *a* associated with one parent/subspecies, hence we report *d*/|*a*| such that it can be interpreted with respect to bacterial abundance. For the vast majority of loci (83.53%), the allele associated with lower abundance is dominant or partially dominant (−1.25 < *d*/|*a*| < −0.75; Fig. 3B). On the basis of the arbitrary cutoffs we used to classify dominance, only a small proportion of alleles are underdominant (0.22%; *d*/|*a*| < −1.25) or overdominant (0.15%; *d*/|*a*| > 1.25). However for one-third of the significant SNPs, the heterozygotes display transgressive phenotypes, i.e. mean abundances that are either significantly lower (31% of SNPs)- or higher (2% of SNPs) than those of both homozygous genotypes. Interestingly, the *domesticus* allele was associated with higher bacterial abundance in two-thirds of this subset (33.2% vs 16.3% *musculus* allele; Fig. 3A).

**Figure 3:**
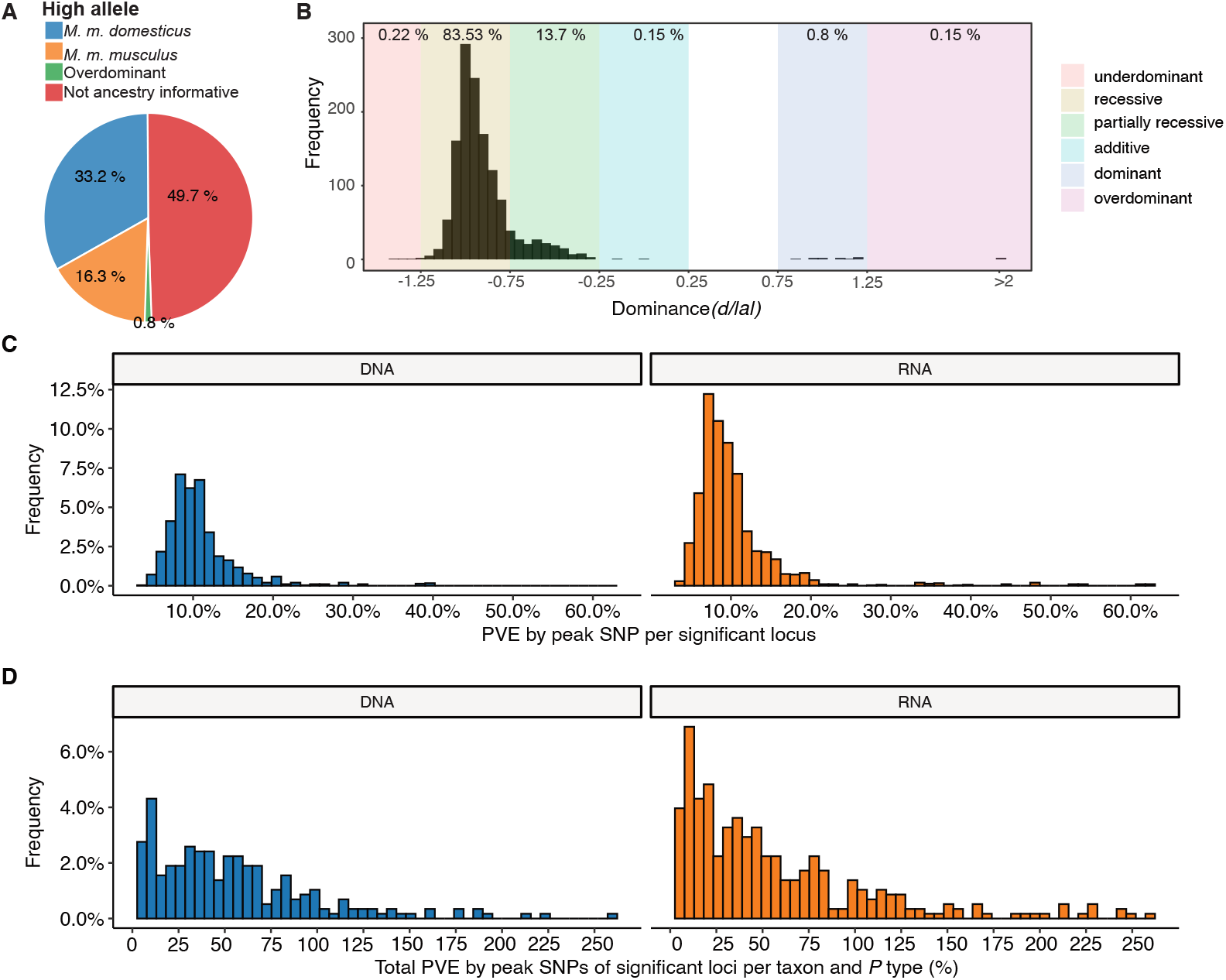
Genetic architecture of significant loci. A) Source of the allele with the highest phenotypic value. B) Histogram of dominance values d/a of significant loci reveals a majority of loci acting recessive or partially recessive. C) Histogram showing the percentage of variance explained (PVE) by the peak SNP for DNA (blue, left) and RNA (orange, right). D) Collective PVE by lead SNPs of significant loci within a taxon. Values are calculated separately for each *P* value type (total, additive, and dominance).

Next, we estimated phenotypic effect sizes by calculating the percentage variance explained (PVE) by the peak SNP of each significant region. Peak SNPs explain between 3% and 64% of the variance in bacterial abundance, with a median effect size of 9.3% (Fig. 3C). The combined effects of all significant loci for each taxon ranged from 4.9% to 259%, with a median of 41.8% (Fig. 3D). Note, combined effects for many taxa (33 out of 59) exceed SNP-heritability estimates (Fig. 1). While exceeding 100% explained variance is biologically possible, as loci can have opposite phenotypic effects, many of these are likely inflated due to the Beavis effect (Beavis, 1994).

### Functional annotation of candidate genes

In order to reveal potential higher level biological phenomena among the identified loci, we performed pathway analysis to identify interactions and functional categories enriched among the genes in significant intervals. We used STRING (Szklarczyk et al., 2019) to calculate a protein-protein interaction (PPI) network of 925 protein-coding genes nearest to significant SNPs (upstream and/or downstream). A total of 768 genes were represented in the STRING database, and the maximal network is highly significant (PPI enrichment *P* value: 2.15 × 10^-14^) displaying 668 nodes connected by 1797 edges and an average node degree of 4.68. After retaining only the edges with the highest confidence (interaction score > 0.9), this results in one large network with 233 nodes, 692 edges and ten smaller networks (Fig. 4).

**Figure 4:**
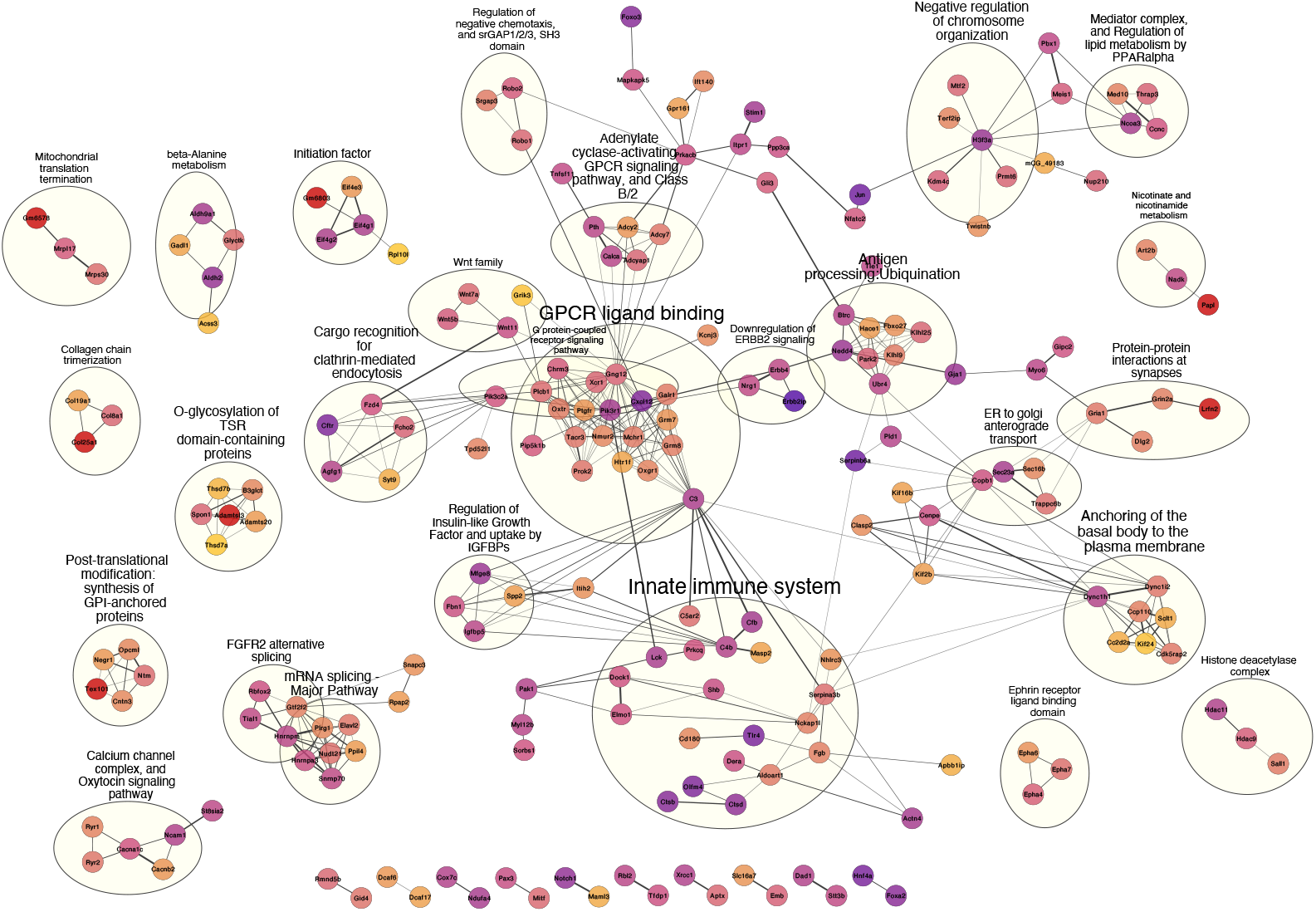
High confidence protein-protein interaction network of genes closest to SNPs significantly associated with bacterial abundances. Network clusters are annotated using STRING’s functional enrichment (Doncheva et al., 2019). Nodes represent proteins and edges their respective interactions. Only edges with an interaction score higher than 0.9 are retained. The width of the edge line expresses the interaction score calculated by STRING. The color of the nodes describe the expression of the protein in the intestine where yellow is not expressed and purple is highly expressed.

Next, we functionally annotated clusters using STRING’s functional enrichment plugin. The genes of the largest cluster are part of the G protein-coupled receptor (GPCR) ligand binding pathway. GPCRs are the largest receptor superfamily and also the largest class of drug targets (Sriram and Insel, 2018). We then calculated the top ten hub proteins from the network based on Maximal Clique Centrality (MCC) algorithm with CytoHubba to predict important nodes that can function as ‘master switches’ (Suppl. Fig. 7). The top ten proteins contributing to the PPI network were GNG12, MCHR1, NMUR2, PROK2, OXTR, XCR1, TACR3, CHRM3, PTGFR, and C3, which are all involved in the GPCR signaling pathway.

Further, we performed enrichment analysis on the 925 genes nearest to significant SNPs using the *clusterprofiler* R package. We found 14 KEGG pathways to be over-represented: circadian entrainment, oxytocin signaling pathway, axon guidance, calcium signaling, cAMP signaling, cortisol synthesis and secretion, cushing syndrome, gastric acid secretion, glutamatergic synapse, mucin type O-glycan biosynthesis, inflammatory mediator regulation of TRP channels, PD-L1 expression and the PD-1 checkpoint pathway in cancer, tight junction, and the *Wnt* signaling pathway (Suppl. Table 5, Suppl. Fig. 8-9). Finally, genes involved in five human diseases are enriched, among them mental disorders (Suppl. Fig. 10).

Finally, due to the observation of a significant enrichment of cospeciating taxa among the bacterial species depleted in early onset IBD (Groussin et al., 2017) and the evidence that IBD is especially associated with a dysbiosis in mucosa-associated communities (Yang et al., 2020a; Daniel et al., 2021), we specifically examined possible over-representation of genes involved in IBD (Khan et al., 2021) among the 925 genes neighboring significant SNPs. We found 14 out of the 289 IBD genes, which was significantly more than expected by chance (10 000 times permuted mean: 2.7, simulated *P* = .0001; Suppl. Table 6). Interestingly, SNPs in five out of the 14 genes are associated with ASVs belonging to the genus *Oscillibacter,* a cospeciating taxon known to decrease during the active state of IBD (Metwaly et al., 2020).

### Comparison of significant loci to published gut microbiome mapping studies

Next, we compiled a list of 648 unique confidence intervals of significant associations with gut bacterial taxa from seven previous mouse QTL studies (Benson et al., 2010; McKnite et al., 2012; Leamy et al., 2014; Wang et al., 2015; Org et al., 2015; Snijders et al., 2016; Kemis et al., 2019) and compared this list to our significance intervals for bacterial taxa at both the DNA and RNA level (346 unique intervals). Regions larger than 10Mb were removed from all studies. We found 434 overlapping intervals, which is significantly more than expected by chance (10 000 times permuted mean: 368, simulated *P*=.0073, see Methods). Several of our smaller significant loci overlapped with larger loci from previous studies and removing this redundancy left 186 significant loci with a median interval size of 0.78 Mb (Fig. 5). The most frequently identified locus is located on chromosome 2 169-171 Mb where protein coding genes *Gm11011, Znf217, Tshz2, Bcas1, Cyp24a1, Pfdn4, 4930470P17Rik*, and *Dok5* are situated.

**Figure 5:**
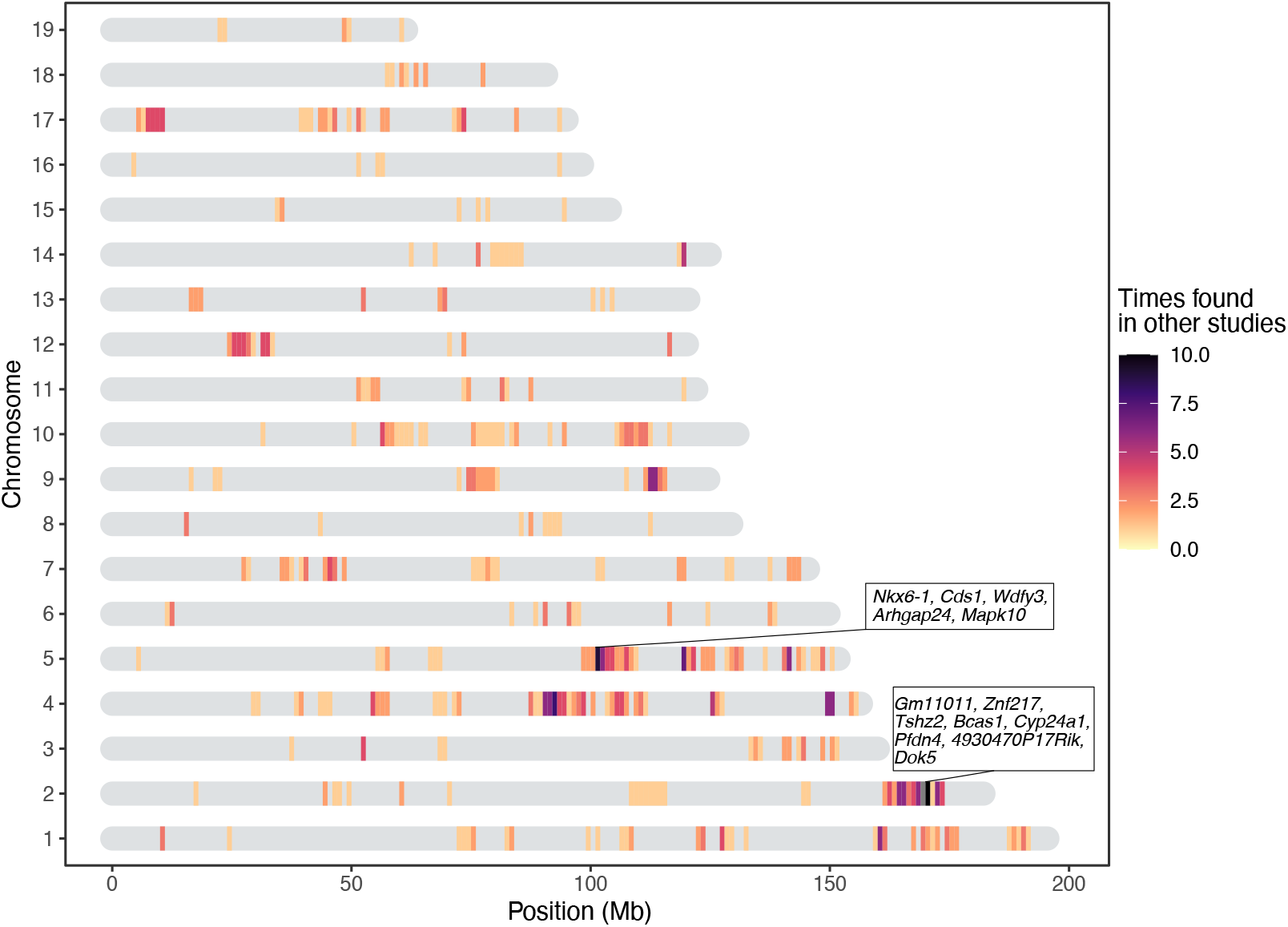
Heatmap showing the significant loci in this study that were previously found in other QTL studies of the mouse gut microbiome. The genes present in two repeatedly identified regions are depicted in boxes.

Additionally, we collected genes within genome-wide significant regions reported in seven human microbiome GWAS (mGWAS) (Bonder et al., 2016; Turpin et al., 2016; Goodrich et al., 2016; Wang et al., 2016; Hughes et al., 2020; Rühlemann et al., 2021; Kurilshikov et al., 2021). However, no significant over-representation of genes was found within our significance intervals (*P* = *.156*), nor within our list of genes closest to a significant SNP (*P* = .62).

### Proteins differentially expressed in germ-free vs conventional mice

To further validate our results, we compared the list of genes contained within intervals of our study to a list of differentially expressed protein between germ-free and conventionally raised mice (Mills et al., 2020). This comparison was made based on the general expectation that genes associated with variation in microbial abundances would be more likely to differ according to the colonization status of the host. Thus, we examined the intersection between genes identified in our study and the proteins identified as highly associated ( |π| >1) with the colonization state of the colon and the small intestine (Mills et al., 2020). Out of the 373 over- or under-expressed proteins according to colonization status, we find 198 of their coding genes to be among our significant loci, of which 17 are the closest genes to a significant marker (*Iyd, Nln, Slc26a3, Slc3a1, Myom2, Nebl, Tent5a, Fxr1, Cbr3, Chrodc1, Nucb2, Arhgef10l, Sucla2, Enpep, Prkcq, Aacs,* and *Cox7c*). This is significantly more than expected by chance (simulated *P*=.0156, 10 000 permutations). Further, analyzing the protein-protein interactions with STRING results in a significant network (*P*=1.73 × 10^-14^, and average node degree 2.4, Suppl. Fig. 11), with *Cyp2c65, Cyp2c55, Cyp2b10, Gpx2, Cth, Eif3k, Eif1, Sucla2,* and *Rpl17* identified as hub genes (Suppl. Fig. 12).

Subsequently, we merged the information from Mills et al. (2020) and the seven previous QTL mapping studies discussed above to further narrow down the most promising candidate genes, and found 30 genes overlapping with our study. Of these 30 genes, six are the closest gene to a significant SNP. These genes are myomesine 2 (*Myom2*), solute carrier family 3 member 1 (*Slc3a1*), solute carrier family 26 member 3 (*Slc26a3*), nebulette (*Nebl*), carbonyl reductase 3 (*Cbr3*), and acetoacetyl-coA synthetase (*Aacs*).

### Candidate genes influencing bacterial abundance

Finally, all previously mentioned candidate genes were combined in one gene set of 304 genes and compiled in a highly significant PPI network (*P* < 1.0 × 10^-16^, average node degree=4.85, see Methods 4.13). Guided by this network, we filtered out genes situated in the same genomic region and kept the gene with the highest connectivity and supporting information (original network see Suppl. Fig. 13). This gave a resulting gene set of 80 candidate genes (Fig. 6 and Suppl. Table 7). The G protein, GNG12 and the complement component 3 C3, are the proteins with the most edges in the network (30 and 25, respectively), followed by MCHR1, CXCL12, and NMUR2 with each 18 edges. Of these 80 highly connected genes, 66 are associated with bacteria that are either co-speciating (co-speciation rate > 0.5; Groussin et al., 2017) and/or have high heritability (> 0.5) suggesting a functionally important role for these bacterial taxa (Suppl. Table 7).

**Figure 6:**
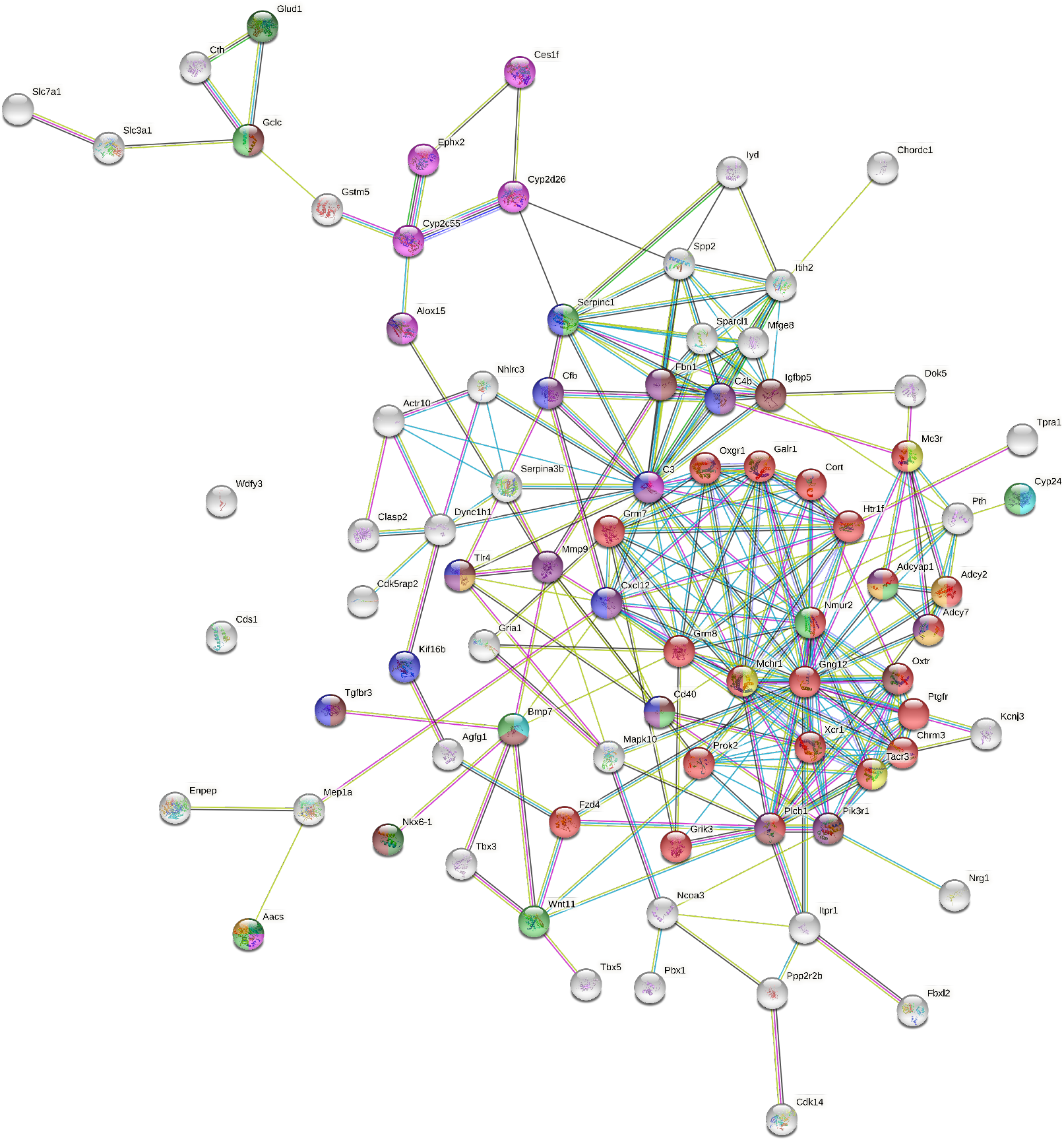
Network of host candidate genes influencing bacterial traits using STRING (https://string-db.org). The nodes represent proteins and are colored according to a selection of enriched GO terms and pathways: G protein coupled receptor (GPCR) signaling (red), regulation of the immune system process (blue), response to nutrient levels (light green), fatty acid metabolic process (pink), glucose homeostasis (purple), response to antibiotic (orange), regulation of feeding behavior (yellow), positive regulation of insulin secretion (dark green), circadian entrainment (brown), and response to vitamin D (turquoise). The color of the edges represents the interaction type: known interactions from curated databases (turquoise) or experimentally determined (pink); predicted interactions from gene neighborhood (green), gene fusions (red), gene co-occurrence (blue); other interactions from textmining (light green), co-expression (black), and protein homology (purple).

## Discussion

Understanding the forces that shape variation in host-associated bacterial communities within host species is key to understanding the evolution and maintenance of meta-organisms. Although numerous studies in mice and humans demonstrate that host genetics influences gut microbiota composition (McKnite et al., 2012; Leamy et al., 2014; Goodrich et al., 2014; Org et al., 2015; Davenport et al., 2015; Wang et al., 2016; Bonder et al., 2016; Goodrich et al., 2016; Kemis et al., 2019; Suzuki et al., 2019; Ishida et al., 2020; Hughes et al., 2020; Rühlemann et al., 2021), our study is unique in a number of important ways. First, the unique genetic resource of mice collected from a naturally occurring hybrid zone together with their native microbes yielded extremely high mapping resolution and the possibility to uncover ongoing evolutionary processes in nature. Second, our study is the first to perform genetic mapping of 16S rRNA transcripts in the gut environment, which was previously shown to be superior to DNA-based profiling in a genetic mapping study of the skin microbiota (Belheouane et al., 2017). Third, our study is one of the only to specifically examine the mucosa-associated community. It was previously reasoned that the mucosal environment may better reflect host genetic variation (Spor et al., 2011), and evidence for this hypothesis exists in nature (Linnenbrink et al., 2013). Finally, by cross-referencing our results with previous mapping studies and recently available proteomic data from germ-free versus conventional mice, we curated a more reliable list of candidate genes and pathways. Taken together, these results provide unique and unprecedented insight into the genetic basis for host-microbe interactions.

Importantly, by using wild-derived hybrid inbred strains to generate our mapping population, we gained insight into the evolutionary association between hosts and their microbiota at the transition from within species variation to between species divergence. Genetic relatedness in our mapping population significantly correlates with microbiome similarity, supporting a basis for codiversification at the early stages of speciation. A substantial proportion of microbial taxa are heritable, and heritability is correlated with cospeciation rates. This suggests that (i) vertical transmission could enable greater host adaptation to bacteria and/or (ii) the greater number of host genes associated with cospeciating taxa could indicate a greater dependency on the host, such that survival outside a specific host is reduced, making horizontal transmission less likely.

By performing 16S rRNA gene profiling at both the DNA and RNA level, we found that 30% (DNA-based) to 45% (RNA-based) of bacterial taxa are heritable, which is consistent with or higher than estimates reported in humans (~10%, Goodrich et al., 2016; ~21%, Turpin et al., 2016) and previous mouse studies (Kovacs et al., 2011; McKnite et al., 2012; Campbell et al., 2012; O’Connor et al., 2014; Carmody et al., 2015; Korach-Rechtman et al., 2019;). The high proportion of heritable taxa, with estimates of up to 91%, is likely explained in part by several factors of our study design. First, mice were raised in a controlled common environment, and heritability estimates in other mammals were shown to be contingent on the environment (Grieneisen et al., 2021). Further, bacterial communities were sampled from cecal tissue instead of fecal content (Linnenbrink et al., 2013), and genetic variation was higher than in a typical mapping study due to subspecies differences. For the RNA-based traits, heritability estimates were significantly correlated with previously reported cospeciation rates in mammals (Groussin et al., 2017). This pattern, as well as the higher proportion of heritable taxa in RNA-based traits, suggest that host genetic effects are more strongly reflected by bacterial activity than cell number.

Accordingly, we found a total of 179 and 313 unique significant loci for DNA-based and RNA-based bacterial abundance, respectively, passing the conservative study-wide significance threshold. Taxa had a median of five significant loci, suggesting a complex and polygenic genetic architecture affecting bacterial abundances. We identify a higher number of loci in comparison to previous QTL and GWAS studies in mice (Benson et al., 2010; McKnite et al., 2012; Leamy et al., 2014; Wang et al., 2015; Org et al., 2015; Snijders et al., 2016; Kemis et al., 2019), which may be due to a number of factors. The parental strains of our study were never subjected to rederivation and subsequent reconstitution of their microbiota, and natural mouse gut microbiota are more variable than the artificial microbiota of laboratory strains (Kohl and Dearing, 2014; Weldon et al., 2015; Suzuki, 2017; Rosshart et al., 2017;). Furthermore, as noted above, our mapping population harbors both within- and between-subspecies genetic variation. We crossed incipient species sharing a common ancestor ~ 0.5 million years ago, hence we may also capture the effects of mutations that fixed rapidly between subspecies due to strong selection, which are typically not variable within species (Walsh, 1998; Barton and Keightley, 2002).

Importantly, our results also help to describe general features of the genetic architecture of bacterial taxon activity. For the majority of loci, the allele associated with lower relative abundance of the bacterial taxon was (partially) dominant. This suggests there is strong purifying selection against a high abundance of any particular taxon, which may help ensure high alpha diversity. The heterozygotes of one-third of significant SNPs displayed transgressive phenotypes. This is consistent with previous studies of hybrids (Turner et al., 2012; Turner and Harr, 2014; Wang et al., 2015;), for example, wild-caught hybrids showed broadly transgressive gut microbiome phenotypes. This pattern can be explained by over- or underdominance, or by epistasis (Rieseberg et al., 1999).

Notably, many loci significantly associated with bacterial abundance in this study were implicated in previous studies (Fig. 5). For example, chromosome 2 169-171 Mb is associated with ASV23 (*Eisenbergiella), Eisenbergiella* and ASV32 (unclassified Lachnospiraceae) in this study, and overlaps with significant loci from three previous studies (Leamy et al., 2014; Snijders et al., 2016; Kemis et al., 2019). This region contains eight protein-coding genes: *Gm11011, Znf217, Tshz2, Bcas1, Cyp24a1, Pfdn4, 4930470P17Rik,* and *Dok5.* Another hotspot is on chromosome 5 101-103 Mb. This locus is significantly associated with four taxa in this study (Prevotellaceae, *Paraprevotella*, ASV7 genus *Paraprevotella* and *Acetatifactor*) and overlaps with associations for Clostridiales, Clostridiaceae, Lachnospiraceae, and Deferribacteriaceae (Snijders et al., 2016). Proteincoding genes in this region are: *Nkx6-1, Cds1, Wdfy3, Arhgap24,* and *Mapk10.* As previous studies were based on rederived mouse strains, identifying significant overlap in the identification of host loci suggests that some of the same genes and/or mechanisms influencing major members of gut microbial communities are conserved even in the face of community ‘reset’ in the context of re-derivation. The identity of the taxa is however not always the same, which suggests that functional redundancy may contribute to these observations, if *e.g.* several bacterial taxa fulfill the same function within the gut microbiome (Moya and Ferrer, 2016; Tian et al., 2020). Additionally, there is significant overlap of genes within loci identified in the current study and proteins differentially expressed in the intestine of germ-free mice compared to conventionally raised mice (Mills et al., 2020). Finally, by analyzing the functions of the genes closest to significant SNPs, we found that 12 of the 14 significantly enriched KEGG pathways were shown to be related to interactions with bacteria (Fonken et al., 2010; Thaiss et al., 2014; Neumann et al., 2014; Thaiss et al., 2015a; Thaiss et al., 2015b; Castoldi et al., 2015; Erdman and Poutahidis, 2016; Thaiss et al., 2016; Deaver et al., 2018; Wu et al., 2018; Peng et al., 2020; Nagpal et al., 2020; Hollander and Kaunitz, 2020; Suppl. Table 5).

To improve the robustness of our results, we combined multiple lines of evidence to prioritize candidates, resulting in a network of 80 genes (Suppl. Table 7). At the center of this network is a set of 22 proteins involved in G-protein coupled receptor signaling (Fig. 6, red nodes). MCHR1, NMUR2, and TACR3 (Fig. 6, yellow) are known to regulate feeding behavior (Saito et al., 1999; Cardoso et al., 2012; Smith et al., 2019), and CHRM3 to control digestion (Gautam et al., 2006; Tanahashi et al., 2009). Gut microbes can produce GPCR agonists to elicit host cellular responses (Cohen et al., 2017; Colosimo et al., 2019; Chen et al., 2019; Pandey et al., 2019). Thus, GPCRs may be key modulators of communication between the gut microbiota and host. Another interesting group of genes are those responding to nutrient levels (*Bmp7, Cd40, Aacs, Gclc, Nmur2, Cyp24a1, Adcyap1, Serpinc1,* and *Wnt11*) (Sethi and Vidal-Puig, 2008; Peier et al., 2009; Townsend et al., 2012; Yi and Bishop, 2015; Shi and Tu, 2015; Toderici et al., 2016; Yasuda et al., 2021; Gastelum et al., 2021;), as gut microbiota affect host nutrient uptake (Chung et al., 2018). In addition, CYP24A1, BMP7 and CD40 respond to vitamin D. Previous studies identified vitamin D/the vitamin D receptor to play a role in modulating the gut microbiota (Wang et al., 2016; Malaguarnera, 2020; Yang et al., 2020b; Singh et al., 2020), and CD40 is known to induce a vitamin D dependent antimicrobial response through IFN-γ activation (Klug-Micu et al., 2013).

Another important category of candidate genes are those involved in immunity. Our most significant SNP was situated downstream of the *Tlr4* gene and was associated with the genus *Dorea* and several *Dorea* species. *Dorea* is a known short chain fatty acid producer (Taras et al., 2002; Reichardt et al., 2018) and interacts with tight junction proteins *Claudin-2* and *Occludin* (Alhasson et al., 2017). *Tlr4* is a member of the Toll-like receptor family, and has been linked with obesity, inflammation, and changes in the gut microbiota (Velloso et al., 2015). These combined results reflect an important role for *Dorea* in fatty acid harvesting and intestinal barrier integrity, both of which could act systemically to activate TLR4 and to promote metabolic inflammation (Cani et al., 2008; Delzenne et al., 2011; Nicholson et al., 2012). Moreover, the SNP with the second lowest *P* value was associated with the same taxa and situated 181 kb upstream of *Irak4.* IRAK4 is rapidly recruited after TLR4 activation to enable downstream activation of the NFκB immune pathway. *Irak4* has previously been associated with a change in bacterial abundance using inbred mice (McKnite et al., 2012; Org et al., 2015).

Finally, we identified notable links between candidate genes and five human diseases (mental disorders, blood pressure finding, systemic arterial pressure, substance-related disorders, and atrial septal deficits; Suppl. Fig. 10). The connection to mental disorders is intriguing as involvement of the gut microbiota is suspected (Kelly et al., 2015; Foster et al., 2017; Cox and Weiner, 2018; Chen et al., 2019; Sarkar et al., 2020; Parker et al., 2020; Flux and Lowry, 2020). Taken together with our finding of an enriched set of GPCRs, this highlights the importance of host-microbial interplay along the gut-brain axis. Moreover, we also identify a significant over-representation of IBD genes (Khan et al., 2021) among the 925 genes nearest to significant SNPs (Suppl. Table 6). Interestingly, SNPs in five out of 14 genes are associated with ASVs belonging to the genus *Oscillibacter*, a highly cospeciating taxon known to decrease during the active state of IBD (Metwaly et al., 2020).

In summary, our study provides a number of novel insights into the importance of host genetic variation in shaping the gut microbiome, in particular for cospeciating bacterial taxa. These findings provide an exciting foundation for future studies of the precise mechanisms underlying host-gut microbiota interactions in the mammalian gut and should encourage future genetic mapping studies that extend analyses to the functional metagenomic sequence level.

## Materials and Methods

### Intercross design

We generated a mapping population using partially inbred strains derived from mice captured in the *M. m. musculus - M. m. domesticus* hybrid zone around Freising, Germany in 2008 (Turner et al., 2012). Originally, four breeding stocks were derived from 8-9 ancestors captured from one (FS, HA, TU) or two sampling sites (HO), and maintained with four breeding pairs per generation using the HAN-rotation outbreeding scheme (Rapp, 1972). Eight inbred lines (two per breeding stock) were generated by brother/sister mating of the 8th generation lab-bred mice. We set up the cross when lines were at the 5th-9th generation of brother-sister meeting, with inbreeding coefficients of > 82%.

We first set up eight G1 crosses, each with one predominantly *domesticus* line (FS, HO - hybrid index <50%; see below) and one predominantly *musculus* line (HA, TU - hybrid index >50%); each line was represented as a dam in one cross and sire in another (Suppl. Fig. 14). One line, FS5, had a higher hybrid index than expected, suggesting there was a misidentification during breeding (see genotyping below). Next, we set up G2 crosses in eight combinations (subcrosses), such that each G2 individual has one grandparent from each of the initial four breeding stocks. We included 40 males from each subcross in the mapping population.

This study was performed according to approved animal protocols and institutional guidelines of the Max Planck Institute. Mice were maintained and handled in accordance with FELASA guidelines and German animal welfare law (Tierschutzgesetz § 11, permit from Veterinäramt Kreis Plön: 1401-144/PLÖ-004697).

### Sample collection

Mice were sacrificed at 91 ± 5 days by CO^2^ asphyxiation. We recorded body weight, body length and tail length, and collected ear tissue for genotyping. The caecum was removed and gently separated from its contents through bisection and immersion in RNAlater (Thermo Fisher Scientific, Schwerte, Germany). After overnight storage in RNAlater at 4° C, the RNAlater was removed and tissue stored at −20° C.

### DNA extraction and sequencing

We simultaneously extracted DNA and RNA from caecum tissue samples using Qiagen (Hilden, Germany) Allprep DNA/RNA 96-well kits. We followed the manufacturer’s protocol, with the addition of an initial bead beating step using Lysing matrix E tubes (MP Biomedical, Eschwege) to increase cell lysis. We used caecum tissue because host genetics has a greater influence on the microbiota at this mucosal site than on the lumen contents (Linnenbrink et al., 2013). We performed reverse transcription of RNA with High-Capacity cDNA Transcription Kits from Applied Biosystems (Darmstadt, Germany). We amplified the V1-V2 hypervariable region of the 16S rRNA gene using barcoded primers (27F-338R) with fused MiSeq adapters and heterogeneity spacers following (Rausch et al., 2016) and sequenced amplicons with 250 bp paired-reads on the Illumina MiSeq platform.

### 16S rRNA gene analysis

We assigned sequences to samples by exact matches of MID (multiplex identifier, 10 nt) sequences processed 16S rRNA sequences using the DADA2 pipeline, implemented in the DADA2 R package, version 1.16.0 (Callahan et al., 2016; Callahan, 2016). Briefly, raw sequences were trimmed and quality filtered with the maximum two ‘expected errors’ allowed in a read, paired sequences were merged and chimeras removed. For all downstream analyses, we rarefied samples to 10,000 reads each. Due to the quality filtering, we have phenotyping data for 286 individuals on DNA level, and 320 G2 individuals on RNA level. We classified taxonomy using the Ribosomal Database Project (RDP) training set 16 (Cole et al., 2014). Classifications with low confidence at the genus level (<0.8) were grouped in the arbitrary taxon ‘unclassified_group’.

We used the phyloseq R package (version 1.32.0) to estimate alpha diversity using the Shannon index and Chao1 index, and beta diversity using Bray-Curtis distance (McMurdie and Holmes, 2013). We defined core microbiomes at the DNA- and RNA-level, including taxa present in > 25% of the samples and with median abundance of non-zero values > 0.2% for amplicon sequence variant (ASV) and genus; and >0.5% for family, order, class and phylum.

### Genotyping

We extracted genomic DNA from ear samples using Qiagen Blood and Tissue 96 well kits (Hilden, Germany), according to the manufacturer’s protocol. We sent DNA samples from 26 G0 mice and 320 G2 mice to GeneSeek (Neogen, Lincoln, NE) for genotyping using the Giga Mouse Universal Genotyping Array (GigaMUGA; Morgan et al., 2015), an Illumina Infinium II array containing 141,090 single nucleotide polymorphism (SNP) probes. We quality-filtered genotype data using plink 1.9 (Chang et al., 2015); we removed individuals with call rates <90% and SNPs that were: not bi-allelic, missing in >10% individuals, with minor allele frequency <5%, or Hardy-Weinberg equilibrium exact test *P* values <1e-10. A total of 64,103 SNPs and all but one G2 individual were retained. Prior to mapping, we LD-filtered SNPs with r^2^ >0.9 using a window of 5 SNPs and a step size of 1 SNP. We retain 32,625 SNPs for mapping.

### Hybrid index calculation

For each G0 and G2 mouse, we estimated a hybrid index – defined as the percentage of *M. m. musculus* ancestry. We identified ancestry-informative SNP markers by comparing GigaMUGA data from ten individuals each from two wild-derived outbred stocks of *M. m. musculus* (Kazakhstan and Czech Republic) and two of *M. m. domesticus* (Germany and France) maintained at the Max Planck Institute for Evolutionary Biology (L.M. Turner and B. Payseur, unpublished data). We classified SNPs as ancestry informative if they had a minimum of 10 calls per subspecies, the major allele differed between *musculus* and *domesticus*, and the allele frequency difference between subspecies was > 0.3. A total of 48,361 quality-filtered SNPs from the G0/G2 genotype data were informative, including 8,775 SNPs with fixed differences between subspecies samples.

### Correlation between host relatedness and microbiome structure

To investigate if host relatedness is correlated with individual variation in microbiome composition, we computed a centered relatedness matrix using the 32,625 filtered SNPs with GEMMA (v 0.98.1; Zhou and Stephens, 2012) and microbial composition-based kinship matrix among individuals based on relative bacterial abundances (Chen et al., 2018). The kinship matrix was calculated with the formula:

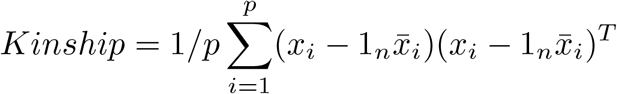

where **X** denotes the *n* × *p* matrix of genotypes or relative abundances, *x_i_* as its *i*th column representing the genotypes of *i*th SNP or the relative abundance of the *i*th ASV, 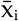 as the sample mean and 1_*n*_ as a *n* × 1 vector of 1’s. We used a Mantel test with the Spearman’s correlation to test for correlation between the host SNP-based kinship and microbial composition-based kinship using 10,000 permutations.

### SNP-based heritability of microbial abundances

We calculated SNP-based heritabilities for bacterial abundances using a linear mixed model implemented in the lme4qtl R package (version 0.2.2; Ziyatdinov et al., 2018). The SNP-based heritability is expressed as:

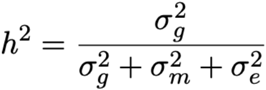

where σ_g_^2^ is the genetic variance estimated by K_SNP_, σ_m_^2^ variance of the mating pair component, and σ_e_^2^ the variance due to residual environmental factors. We determined significance of the heritability estimates using exact likelihood ratio tests, following Supplementary Note 3 in Ziyatdinov et al., 2018, using the exactLRT() function of the R package RLRsim (version 3.1-6; Fabian et al., 2008).

### Genome-wide association mapping

Prior to mapping, we inverse logistic transformed bacterial abundances using the inv.logit function from the R package gtools (version 3.9.2; Gregory R. Warnes, 2020).

We performed association mapping in the R package lme4qtl (version 0.2.2; Ziyatdinov et al., 2018) with the following linear mixed model:

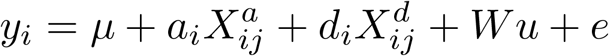

where *y_j_* is the phenotypic value of the *j*th individual; *μ* is the mean, *X_aij_* the additive and *X_dij_* the dominance genotypic index values coded as for individual *j* at locus *i. a* and *d* indicate fixed additive and dominance effects, *W* indicates random effects mating pair and kinship matrix, plus residual error *e*.

We estimated additive and dominance effects separately because we expected to observe underdominance and overdominance in our hybrid mapping population, as well as additive effects, and aimed to estimate their relative importance. To model the additive effect (i.e. 1/2 distance between homozygous means), genotypes at each locus, *i*, were assigned additive index values (X^a^ ∈ 1, 0, −1) for AA, AB, BB, respectively, with A indicating the major allele and B the minor allele. To model dominance effects (i.e. heterozygote mean - midpoint of homozygote means), genotypes were assigned dominance index values (X^d^ ∈ 0, 1) for homozygotes and heterozygotes, respectively.

We included mating pair as a random effect to account for maternal effects and cage effects, as male litter mates are kept together in a cage after weaning. We included kinship coefficient as a random effect in the model to account for population and family structure. To avoid proximal contamination, we used a leave-one-chromosome-out approach, that is, when testing each single-SNP association we used a relatedness matrix omitting markers from the same chromosome (Parker et al., 2014). Hence, for testing SNPs on each chromosome, we calculated a centered relatedness matrix using SNPs from all other chromosomes with GEMMA (v0.97; Zhou and Stephens, 2012). We calculated *P* values for single-SNP associations by comparing the full model to a null model excluding fixed effects. Code for performing the mapping is available at https://github.com/sdoms/mapping_scripts.

We evaluated significance of SNP-trait associations using two thresholds; first, we used a genome-wide threshold for each trait, where we corrected for multiple testing across markers using the Bonferroni method (Abdi, 2007). Second, as bacteria interact with each other within the gut as members of a community, bacterial abundances are non-independent, so we calculated a study-wide threshold dividing the genome-wide threshold by the number of effective taxa included. We used matSpDlite (Nyholt, 2019; Li and Ji, 2005; Qin et al., 2020) to estimate the number of effective bacterial taxa based on eigenvalue variance.

To estimate the genomic interval represented by each significant LD-filtered SNP, we report significant regions defined by the most distant flanking SNPs in the full pre-LD-filtered genotype dataset showing *r^2^* > 0.9 with each significant SNP. We combined significant regions less than 10 Mb apart into a single region. Genes situated in significant regions were retrieved using biomaRt (Steffen Durinck, 2009), and the mm10 mouse genome.

### Dominance analyses

We classified dominance for SNPs with significant associations on the basis of the d/a ratio (Falconer, 1996) where d is the dominance effect, a the additive effect. As the expected value under purely additive effects is 0. As our mapping population is a multi-parental-line cross, and not all SNPs were ancestry-informative with respect to *musculus/ domesticus*, the sign of *a* effects is defined by the major allele within our mapping population, which lacks clear biological interpretation. To provide more meaningful values, we report d/|a|, such that a value of 1 = complete dominance of the allele associated with higher bacterial abundance, and a value of −1 = complete dominance of the allele associated with lower bacterial abundance. Values above 1 or below −1 indicate over/underdominance. We classified effects of significant regions the following arbitrary d/|a| ranges to classify dominance of significant regions (Burke et al., 2002; Miller et al., 2014): underdominant <-1.25, high abundance allele recessive between −1.25 and −0.75, partially recessive between −0.75 and −0.25, additive between −0.25 and 0.25, partially dominant between 0.25 and 0.75, dominant 0.75 and 1.25, and overdominant >1.25.

### Gene ontology and network analysis

The nearest genes up- and downstream of the significant SNPs were identified using the locateVariants() function from the VariantAnnotation R package (version 1.34.0; Valerie et al., 2014) using the default parameters. A maximum of two genes per locus were included (one upstream, and one downstream of a given SNP).

To investigate functions and interactions of candidate genes, we calculated a a protein-protein interaction (PPI) network with STRING version 11 (Szklarczyk et al., 2019), on the basis of a list of the closest genes to all SNPs with significant trait associations. We included network edges with an interaction score >0.9, based on evidence from fusion, neighborhood, co-occurrence, experimental, text-mining, database, and co-expression. We exported this network to Cytoscape v 3.8.2 (Shannon et al., 2003) for identification of highly interconnected regions using the MCODE Cytoscape plugin (Bader and Hogue, 2003), and functional annotation of clusters using the stringApp Cytoscape plugin (Doncheva et al., 2019).

We identified overrepresented KEGG pathways and human diseases using the clusterprofiler R package (version 3.16.1; Yu et al., 2012). *P* values were corrected for multiple testing using the Benjamini-Hochberg method. Pathways and diseases with an adjusted *P* value < .05 were considered over-represented.

### Calculating overlap with other studies and over-representation of IBD genes

To test for significant overlap with loci identified in previous mapping studies and for over-representation of IBD genes, we used the tool *poverlap* (Brent Pedersen, 2013) to compare observed overlap to random expectations based on 10,000 permutations of significant regions. We identified genes within overlapping regions using the locateVariants() function from the VariantAnnotation R package (version 1.34.0; Valerie et al., 2014).

### Combination of results

Hub genes SNP network and their first neighbors, the hub genes from the ‘differentially expressed in GF mice’-network and their respective first neighbors, genes found in both Mills et al. (2020) and other mouse QTL studies, closest genes to a SNP found in Mills et al. (2020), genes situated in the 20 smallest intervals, six genes in the two intervals with the lowest *P* values, twenty genes in intervals found in most different taxa, genes situated in the region with most overlap within our study, and finally the genes situated in the intervals that most frequently overlapped with other studies were combined into on gene set and analyzed with STRING. Genes situated in the same genomic locus were curated according to the number of edges in the STRING network.

## Data and code availability

DNA- and RNA-based 16S rRNA gene sequences are available under project accession number PRJNA759194. Code is available at https://github.com/sdoms/mapping_scripts.

## Supplementary Materials

**Suppl. Fig 1-14, Suppl. Table 1: Heritability estimates, Suppl. Table 2: Genome-wide significant associations, Suppl. Table 3: Study-wide significant associations, Suppl. Table 4: Intervals smaller than 1Mb, Suppl. Table 5: Over-represented KEGG pathways, Suppl. Table 6: IBD genes, Suppl. Table 7: Candidate genes.**

## Acknowledgments

We thank Diethard Tautz for generous support of mouse breeding and Camilo Medina and the MPI-Plön mouse team for performing mouse husbandry, and Katja Cloppenborg-Schmidt and Dr. Sven Künzel for their excellent technical assistance. We thank Mathieu Groussin for assistance with cospeciation rate data. Research funding for this project was provided by the Deutsche Forschungsgemeinschaft Collaborative Research Center 1182, Origin and Function of Metaorganisms’ (J.F.B. and A.F.) and TU 500/2-1 to L.M.T, and by the Max Planck Society (to D. Tautz).

## Author contributions

Conceptualization: L.M.T., S.I., A.F., and J.F.B; Methodology: L. M.T, J.F.B., S.D., S.I., A.F., and A.K.; Software: S.D., M.R., and L.M.T; Validation: S.D., M. R., A.K., and L.M.T.; Formal Analysis: S.D., M.R., A.K., and L.M.T; Investigation: S.D. and L.M.T.; Resources: S.D., H.F., and C.C; Writing - Original Draft: S.D.; Writing - Review & Editing: S.D., L.M.T, and J.F.B; Visualization: S.D. and L.M.T; Supervision: L.M.T., A.F., and J.F.B.; Project Administration: J.F.B.; Funding Acquisition: A.F., L.M.T, and J.F.B.

## Conflicts of Interest

The authors declare no conflict of interest.

## Supplementary figures

**Supplementary figure 1:**
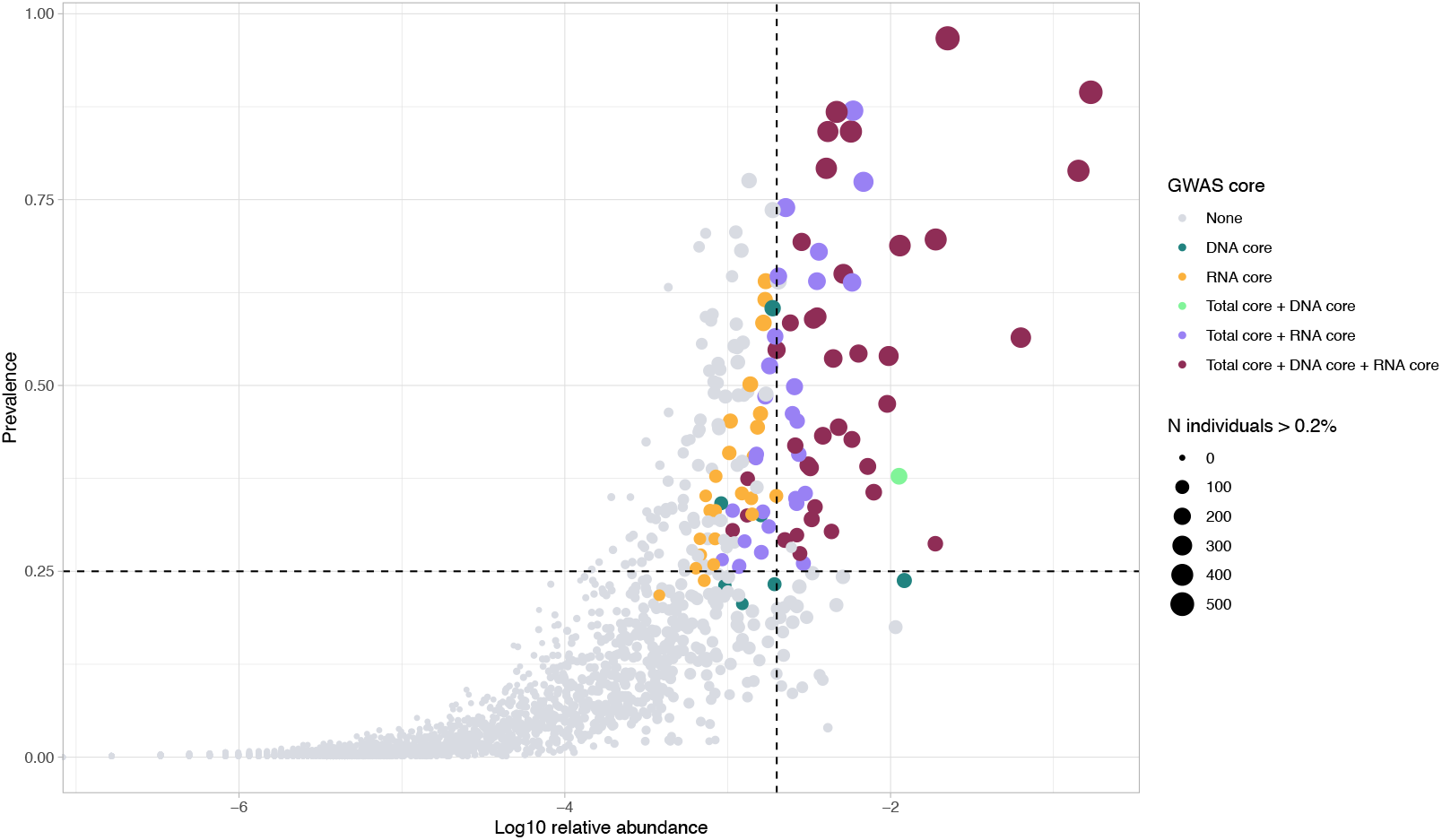
Selection of taxa for mGWAS analysis. A scatter plot showing the association of average relative abundance of taxa with their prevalence in the G2 mapping population. Taxa retained for analysis are colored according to the originating core. The size of each dot represents the number of individuals that have a median abundance higher than 0.2% of the taxon. The dashed lines represent the thresholds of the core (vertical: median abundance>0.2% and horizontal prevalence of 25 %.

**Supplementary figure 2:**
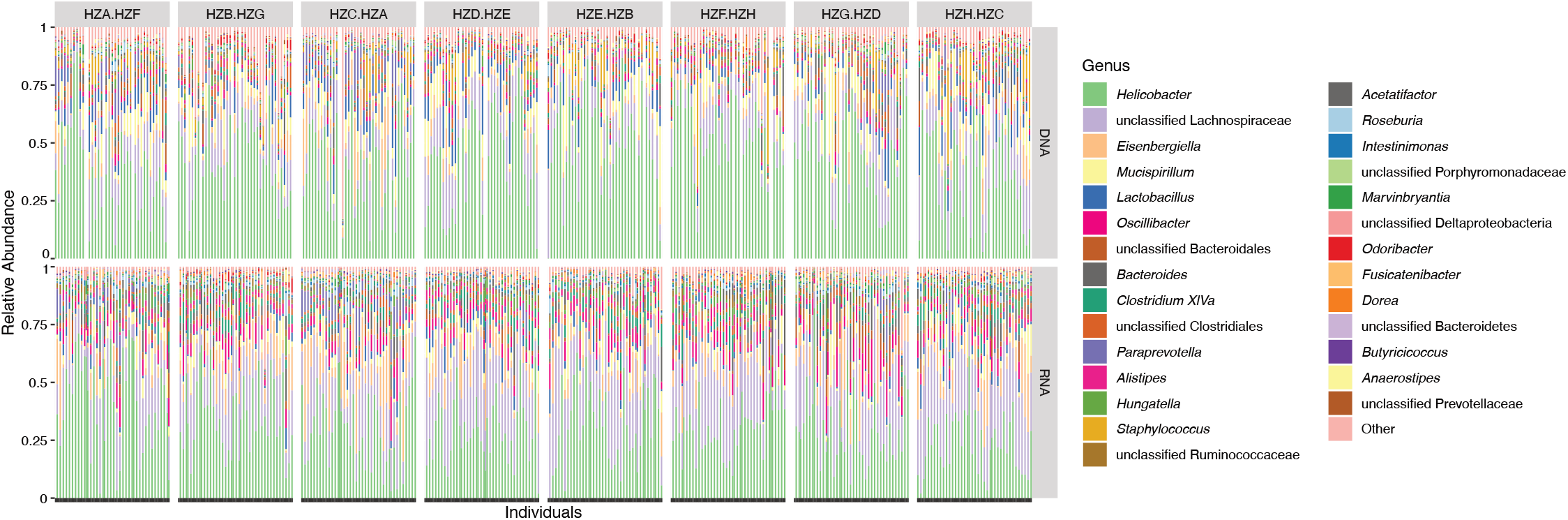
Relative abundances of core genera in G2 mapping population. Each vertical line represents one individual. Subcross (see supplementary figure 14) is indicated at the top.

**Supplementary figure 3:**
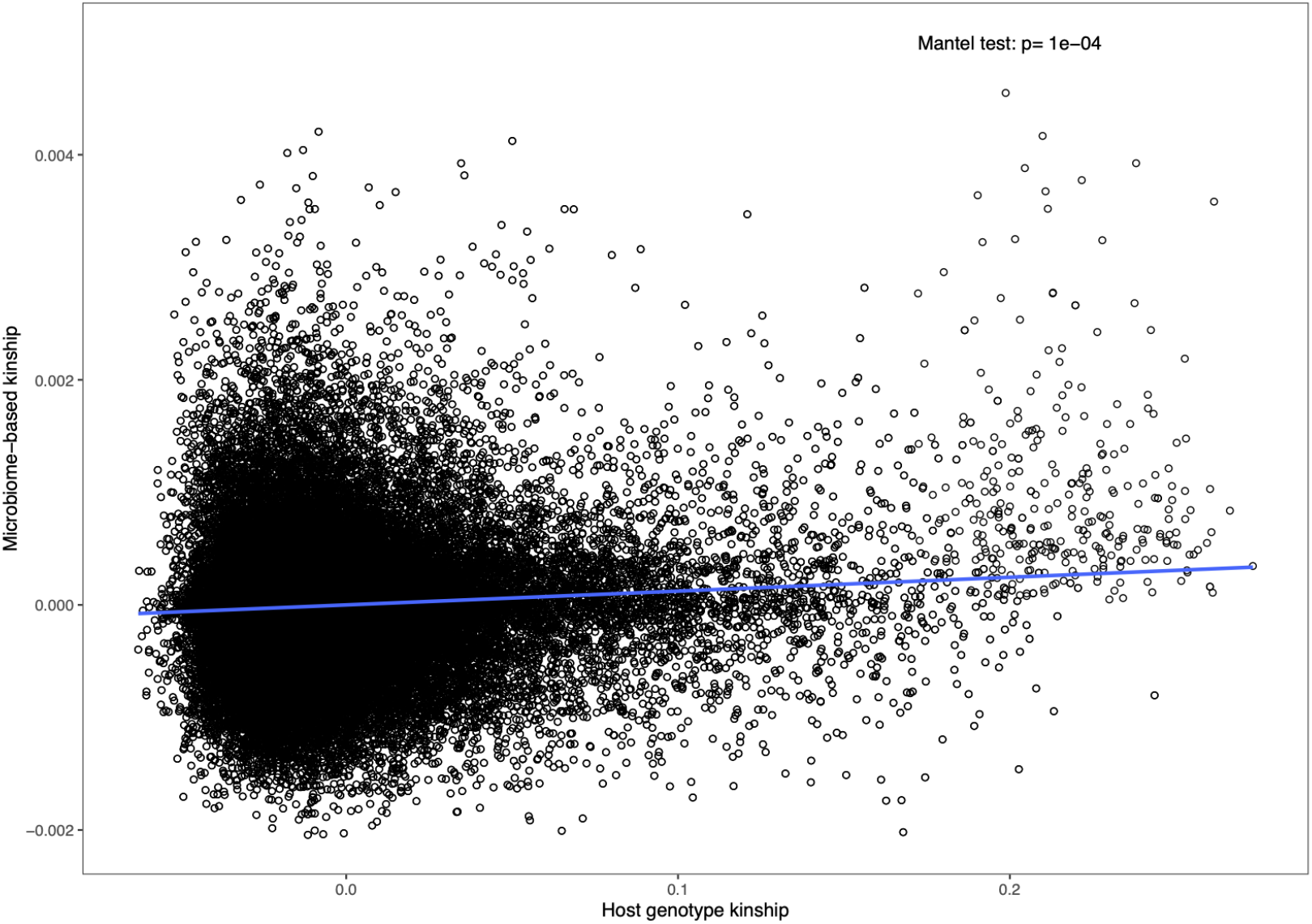
Host genetic relatedness calculated from SNP data (x-axis) correlated with microbial composition-based relatedness (y-axis) calculated from ASV abundances. The blue line represents a linear regression fit to the data.

**Supplementary figure 4:**
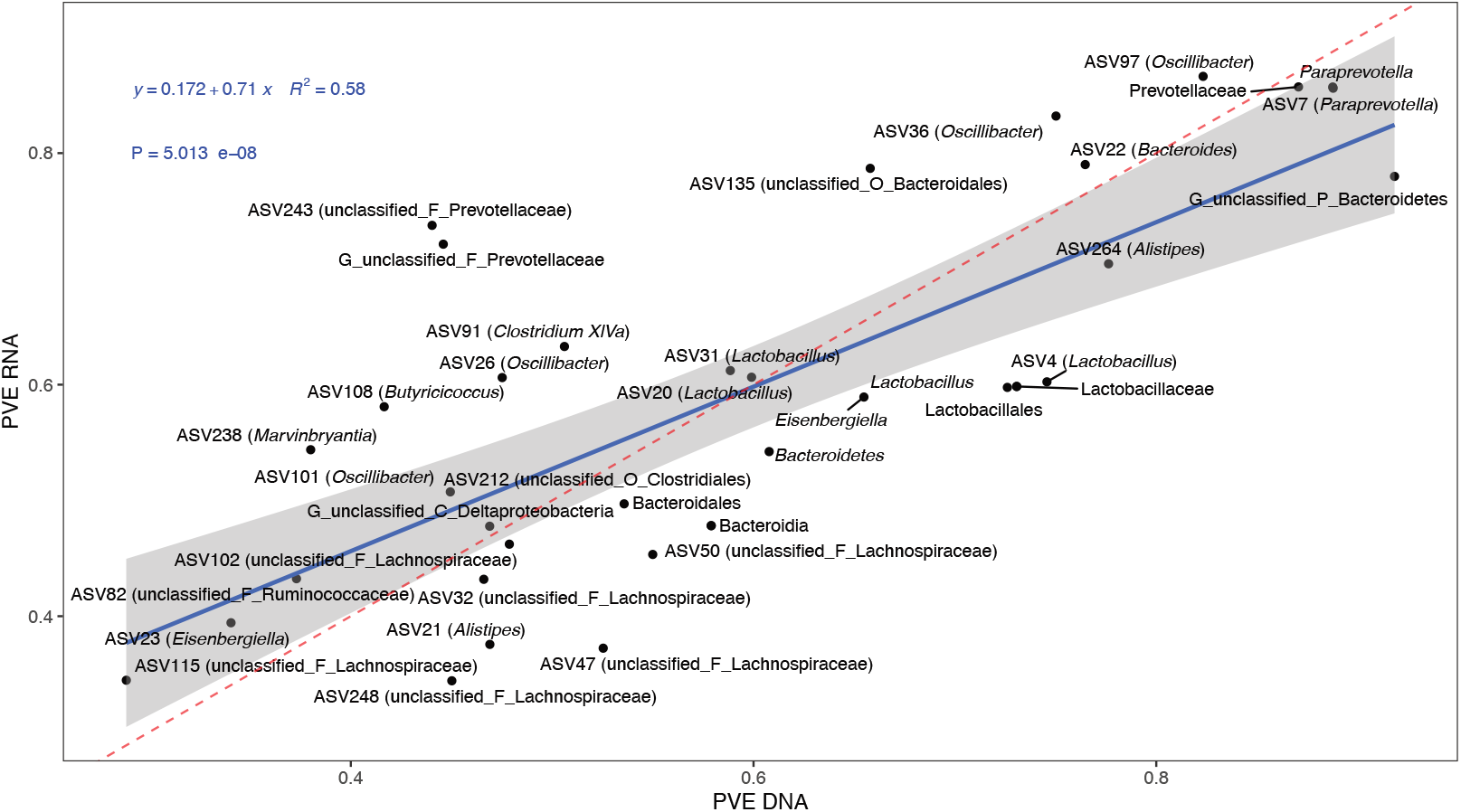
Correlation of SNP-based heritability estimates based on DNA (x-axis) or RNA (y-axis). The blue line represents a linear regression fit to the data. Red dashed line represents the identity line with a slope of 1.

**Supplementary figure 5:**
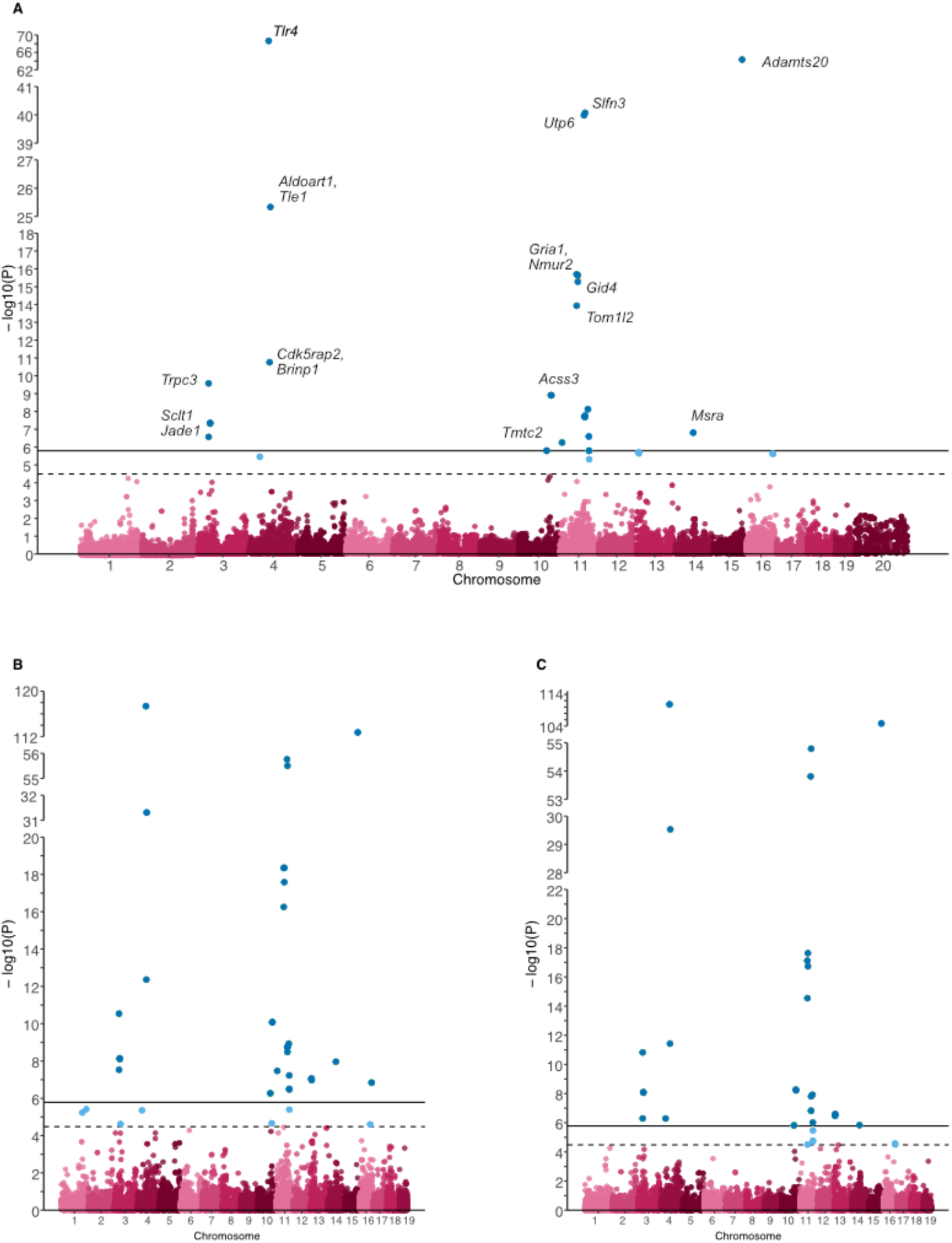
Manhattan plots for ASV184 (*Dorea*) of the complete model (A), the additive effect (B) or the dominance effect (C). SNPs passing the study-wide significance threshold (solid line) are shown in dark blue, while genome-wide significant SNPs (dashed line) are shown in light blue. In panel A, the closest gene to the SNP is shown for a subset of significant SNPs.

**Supplementary figure 6:**
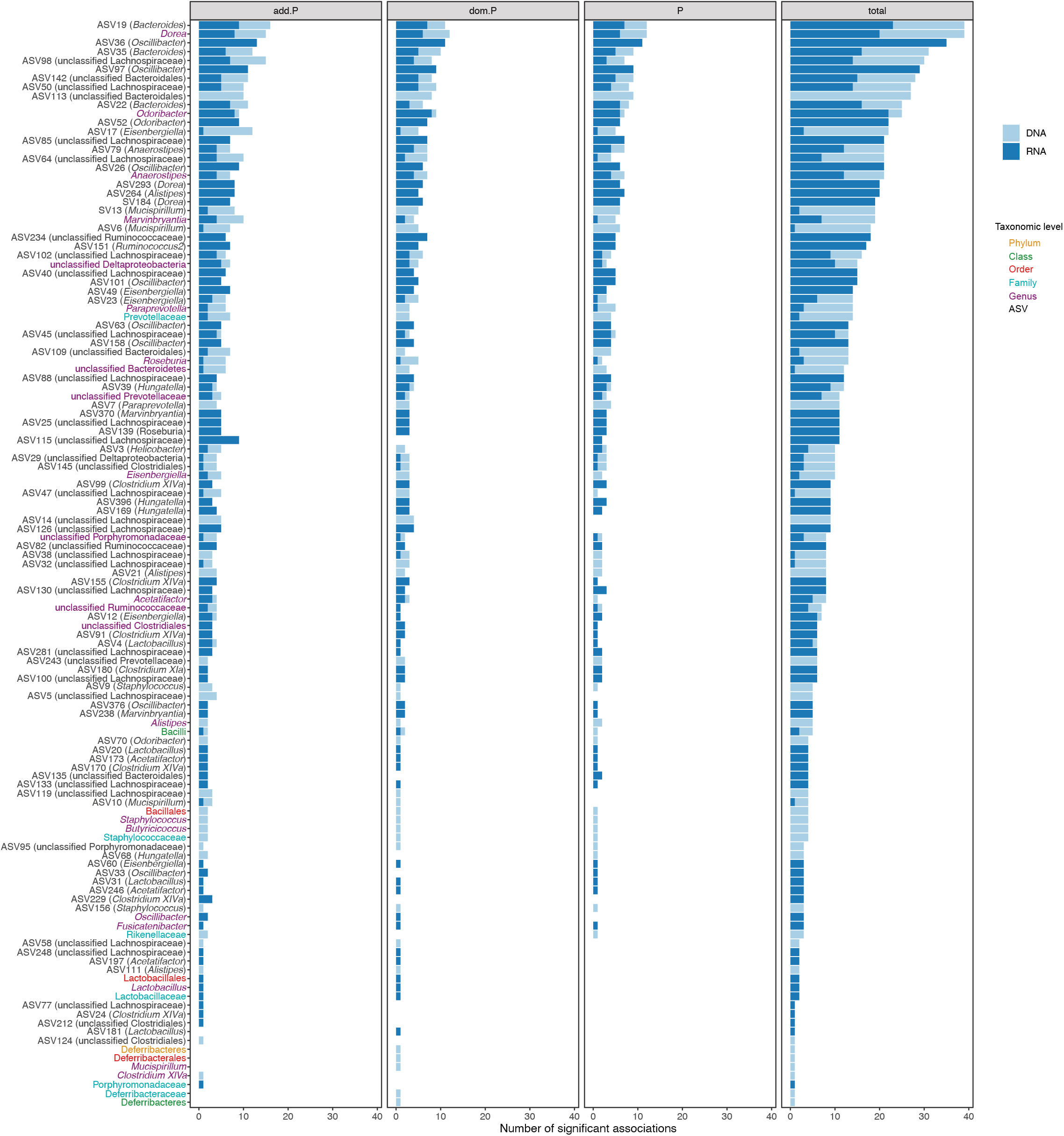
Number of significantly associated loci per bacterial taxon. Loci with significant additive effects (add.P), dominance effects (dom.P) or effects in full model (P) are indicated.

**Supplementary figure 7:**
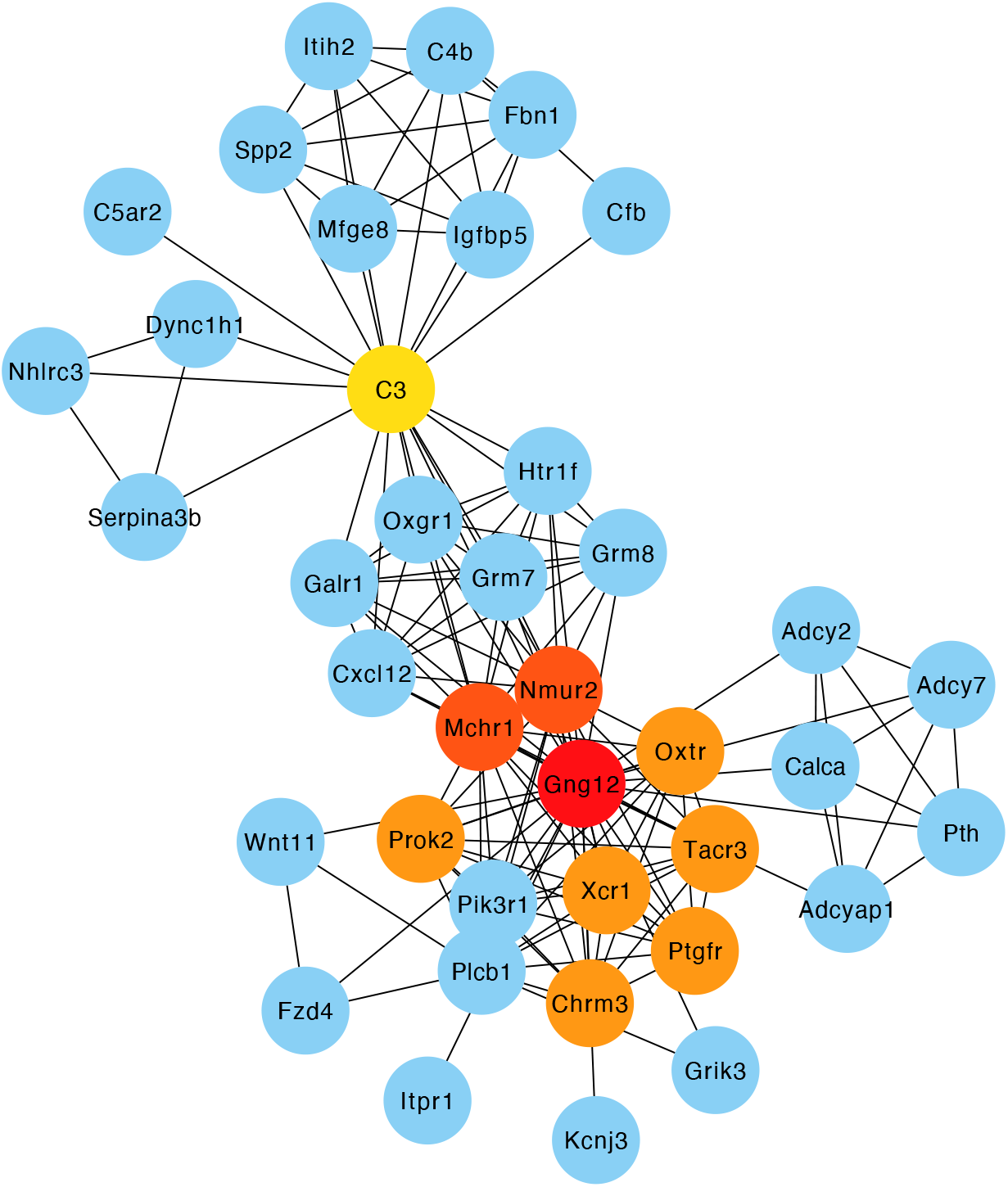
Top ten hub genes of the protein-protein interaction (PPI) network with the closest genes to the host SNPs significantly associated with bacterial abundances. The nodes are colored according to hub gene rank from 1 (red) to 10 (yellow). Blue nodes are the first neighbors.

**Supplementary figure 8:**
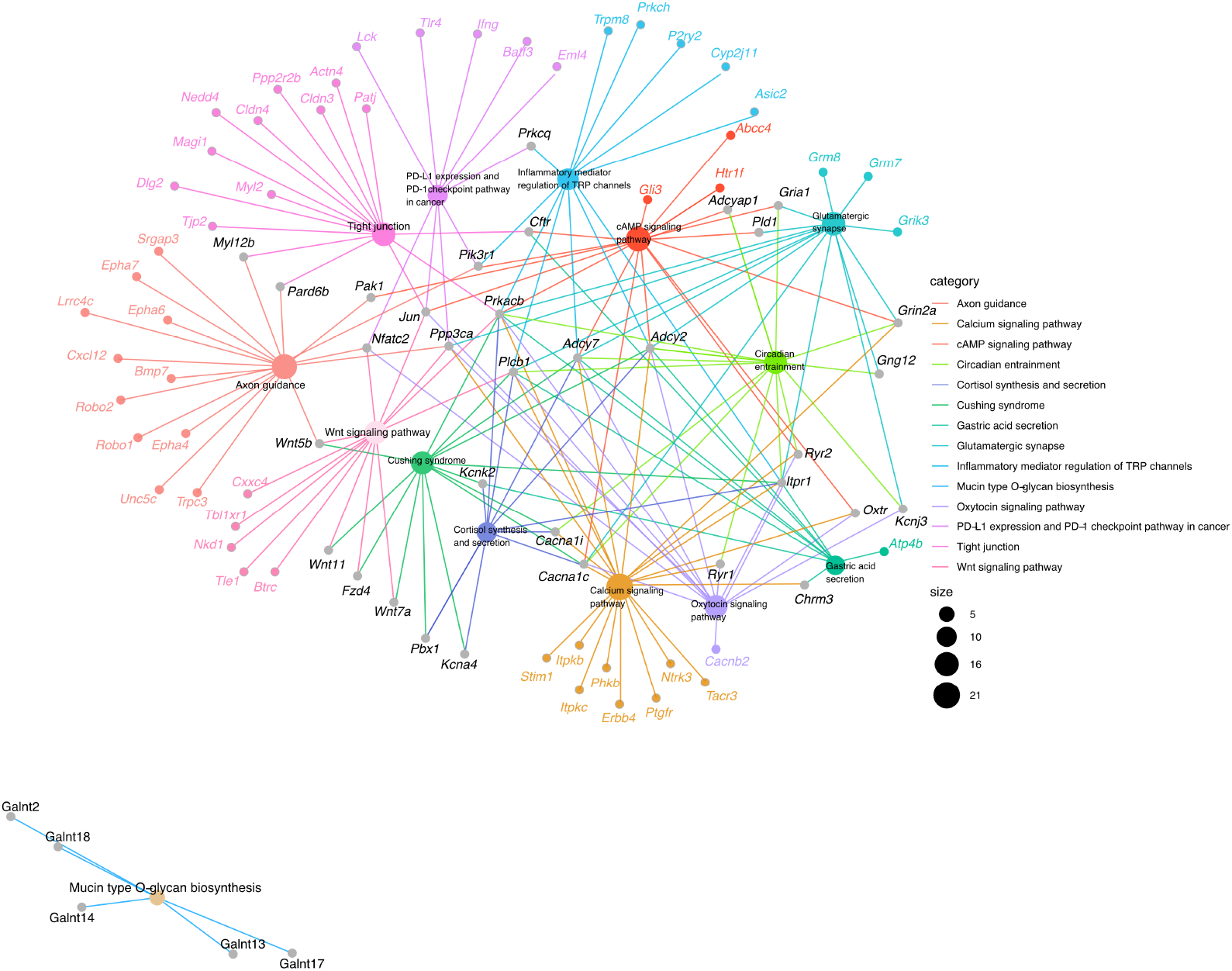
Genes belonging to over-represented KEGG pathways within the host genes closest to significant SNPs from association analysis.

**Supplementary figure 9:**
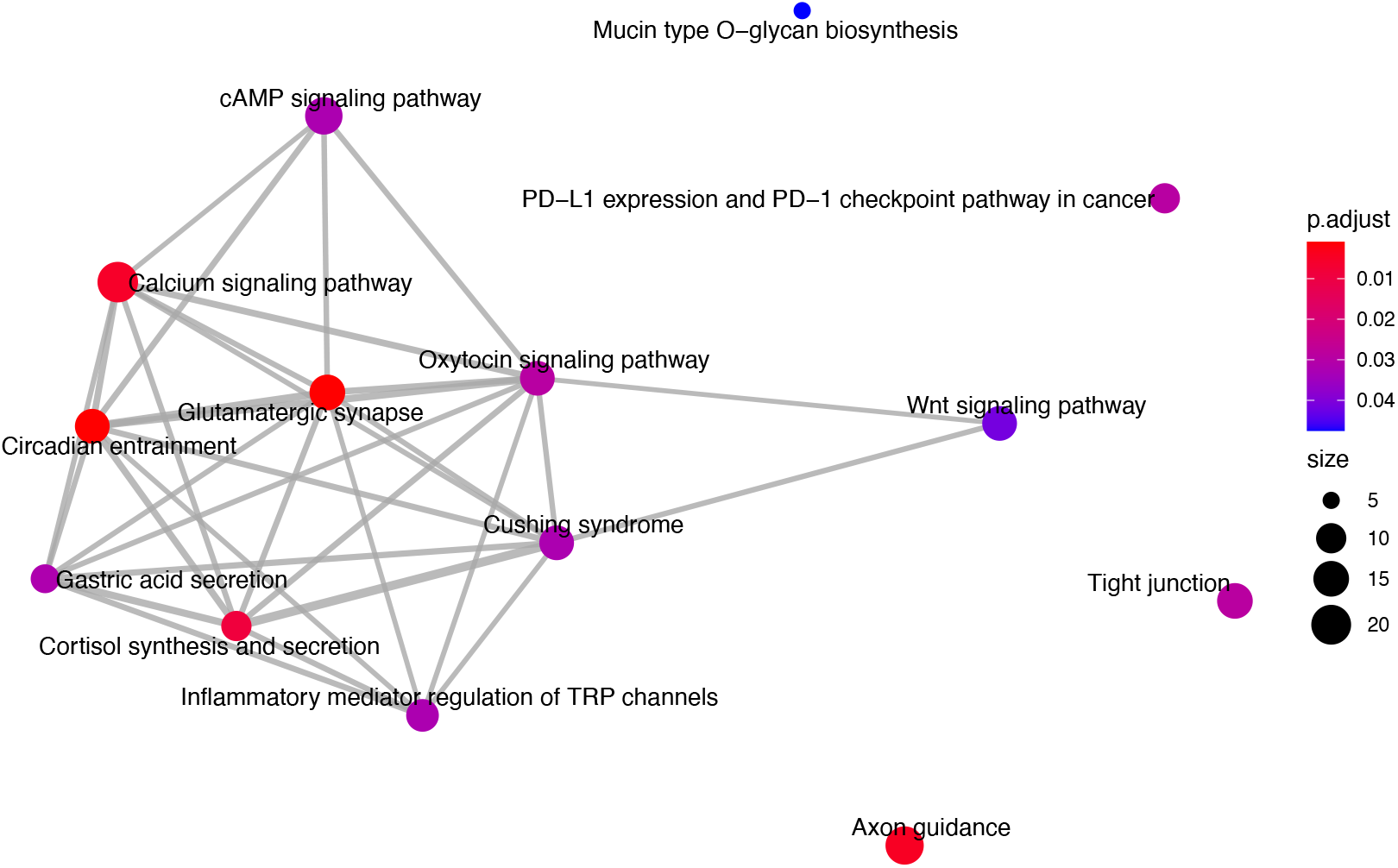
Enriched KEGG pathways among closest genes to significant SNPs from association analysis. Node color indicates FDR-adjusted *P* value of enrichment and node size indicates number of candidate genes in pathway.

**Supplementary figure 10:**
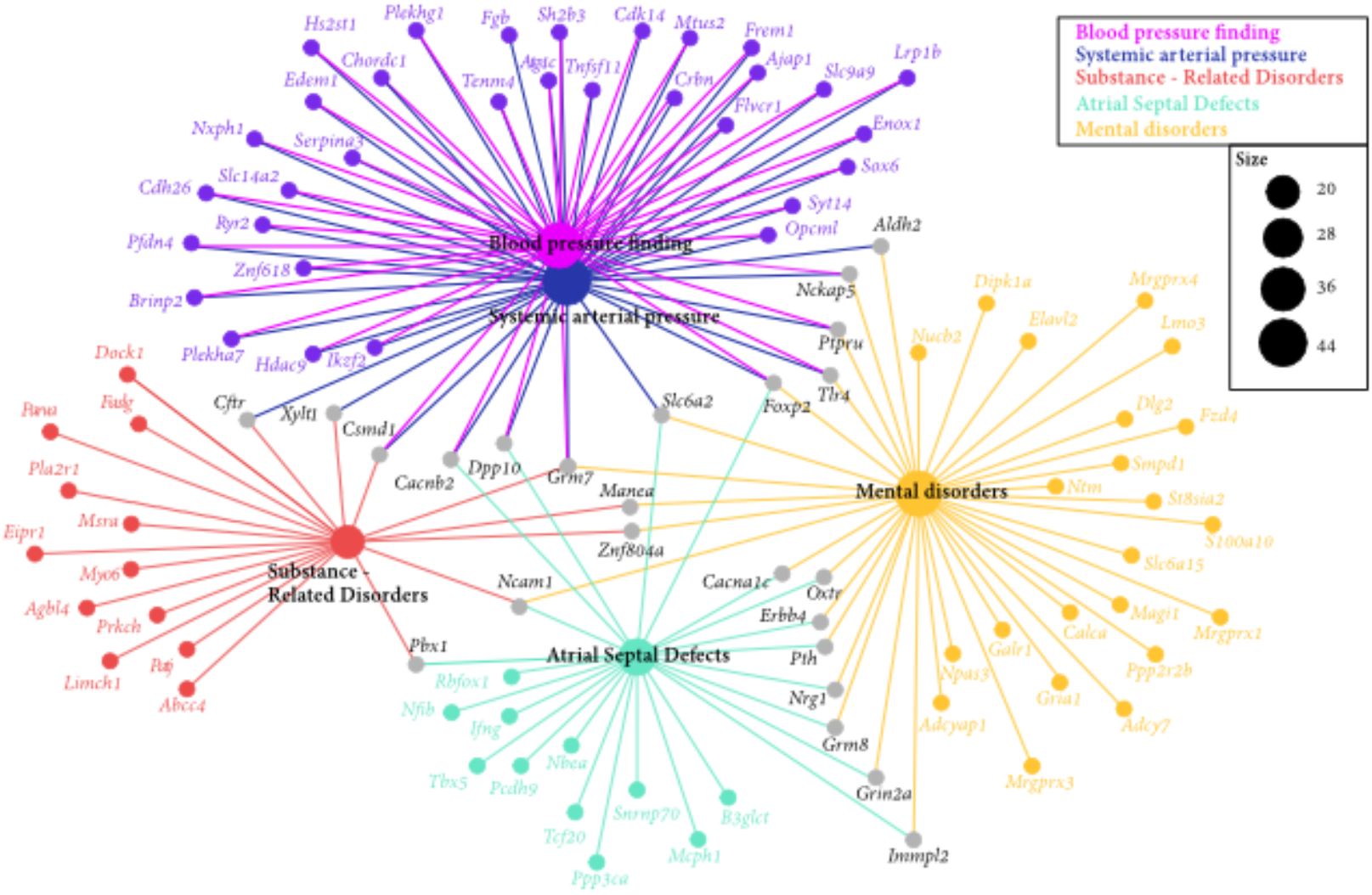
Enriched human diseases among genes closest to significant SNPs from association analysis.

**Supplementary figure 11:**
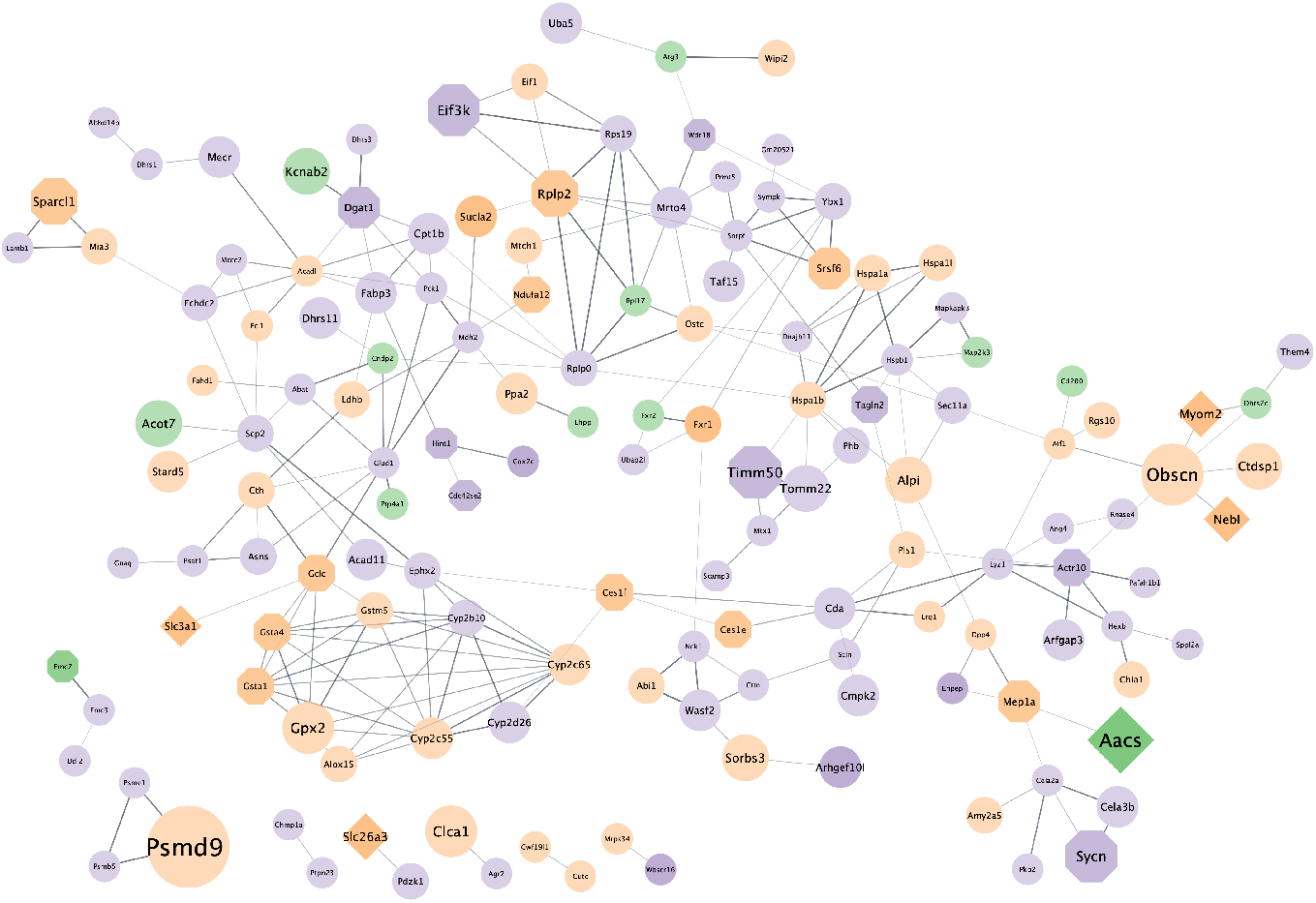
STRING (Szklarczyk et al., 2019) protein-protein interaction network of proteins that are differentially expressed in the intestine (small intestine and colon) of germ-free (GF) mice compared to conventionally raised mice, found in the present study. The color of the network nodes indicates whether the QTL hit was found using the DNA abundances (green), RNA abundances (purple) or was found in both (orange). The shape represents if the gene of the protein was the closest gene to the significant SNP (rectangle), if the gene was also found in QTLs of other studies (octagon), a combination of both (diamond), or only differentially expressed in GF mice vs. conventionally raised mice. The node size expresses the number of taxa where the gene was found in a QTL. The edges represent protein-protein interactions, where the line thickness indicates the strength of the data support from text mining, experiments, databases, co-expression, gene-fusion, and co-occurrence.

**Supplementary figure 12:**
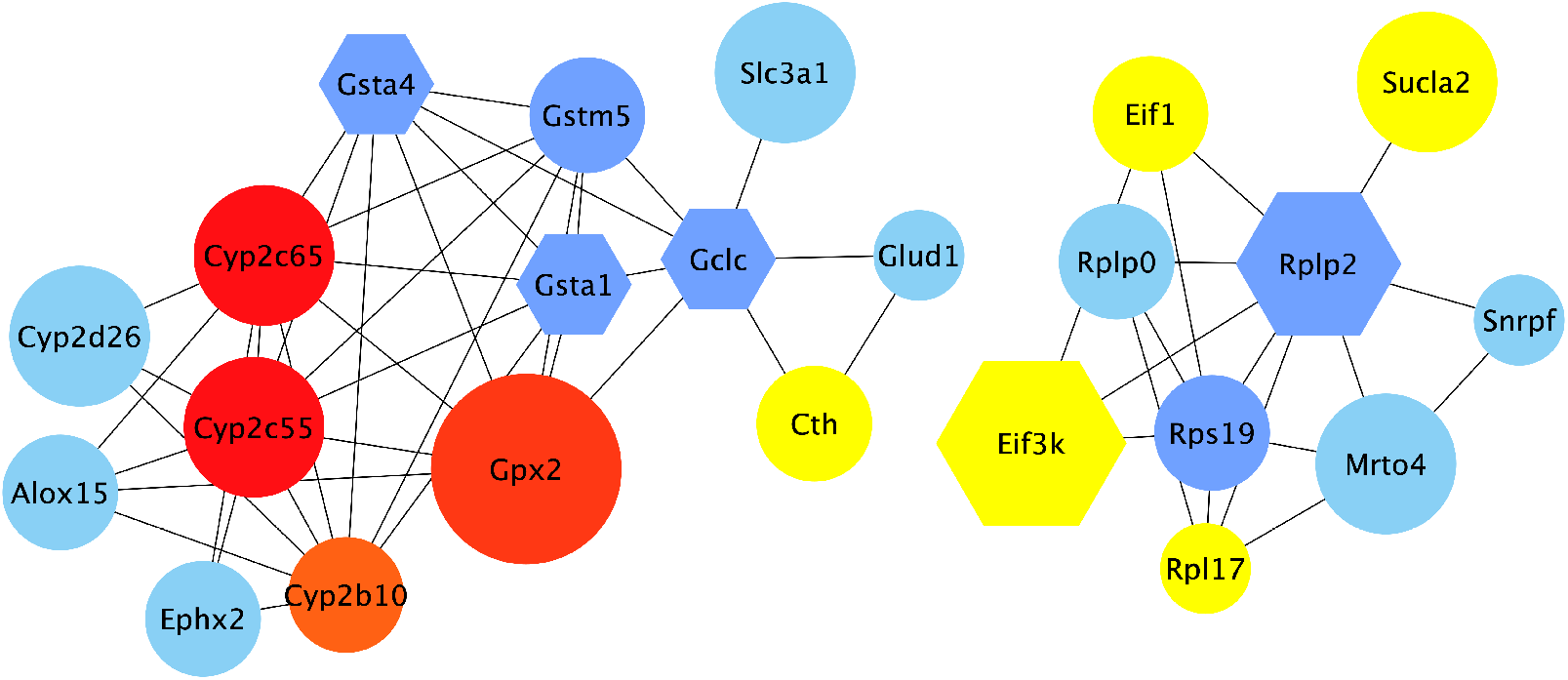
Visualization of the top hub genes calculated with the MCC algo-rithm and their first neighbors from the protein-protein interaction (PPI) network of genes found in intervals in present study that are also differentially expressed in germ-free versus conventionally raised mice. Edges represent the protein-protein associations. The red nodes represent genes with a high degree (= hub genes), and the yellow nodes with a low degree, while the blue nodes represent their first neighbors. All nodes shown are differentially expressed in GF mice. Hexagon shaped nodes are genes/proteins also found associated with gut microbiome abundances in other mouse QTL studies, and round nodes are ‘only’ differentially expressed in GF mice. The size of the node is an indication of the amount of taxa associated with the gene.

**Supplementary figure 13:**
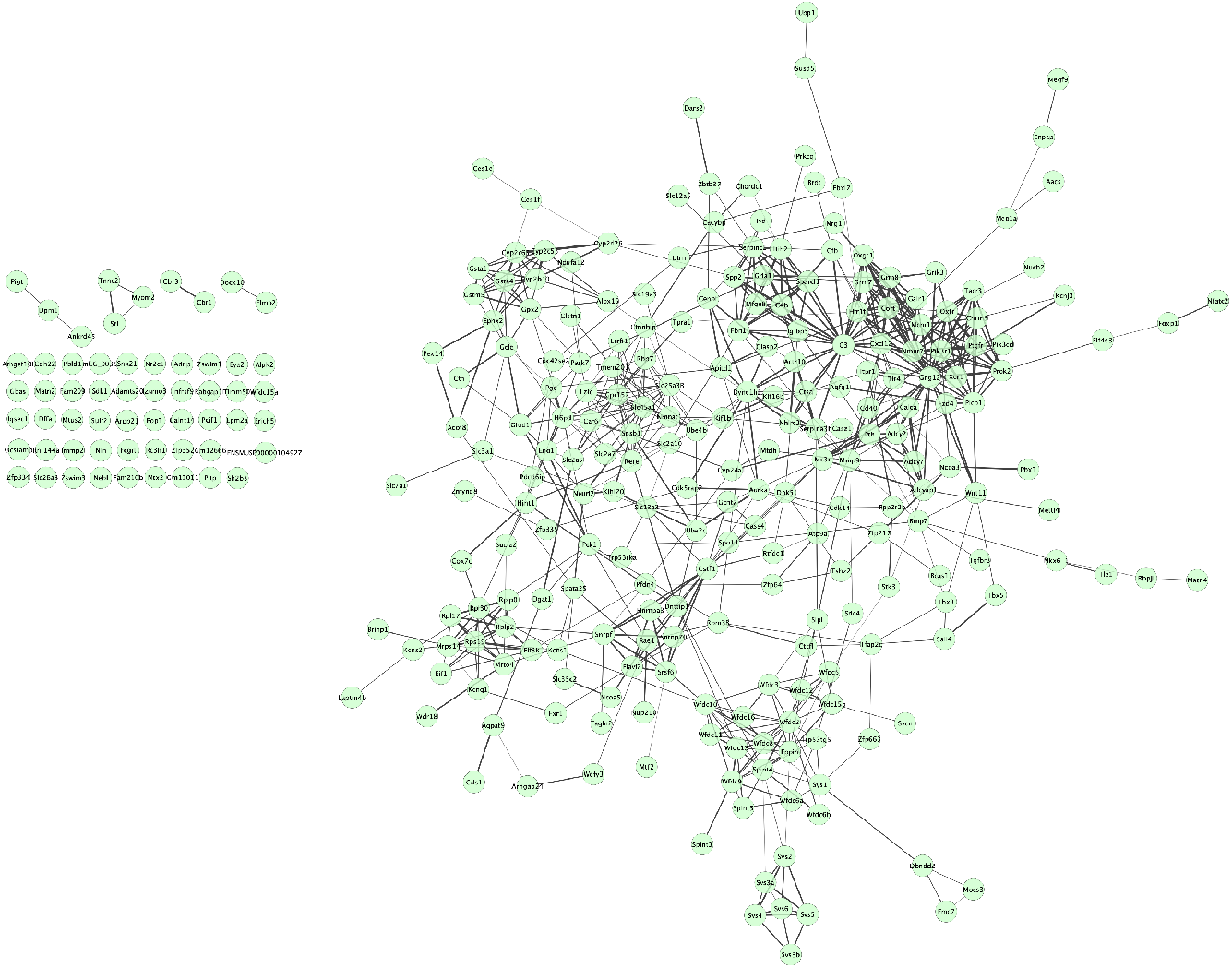
Original protein protein interaction (PPI) network of 304 candidate genes closest to SNPs significantly associated with bacterial abundances. Generated in STRING (Szklarczyk et al., 2019) and Cytoscape (Shannon et al., 2003).

**Supplementary figure 14:**
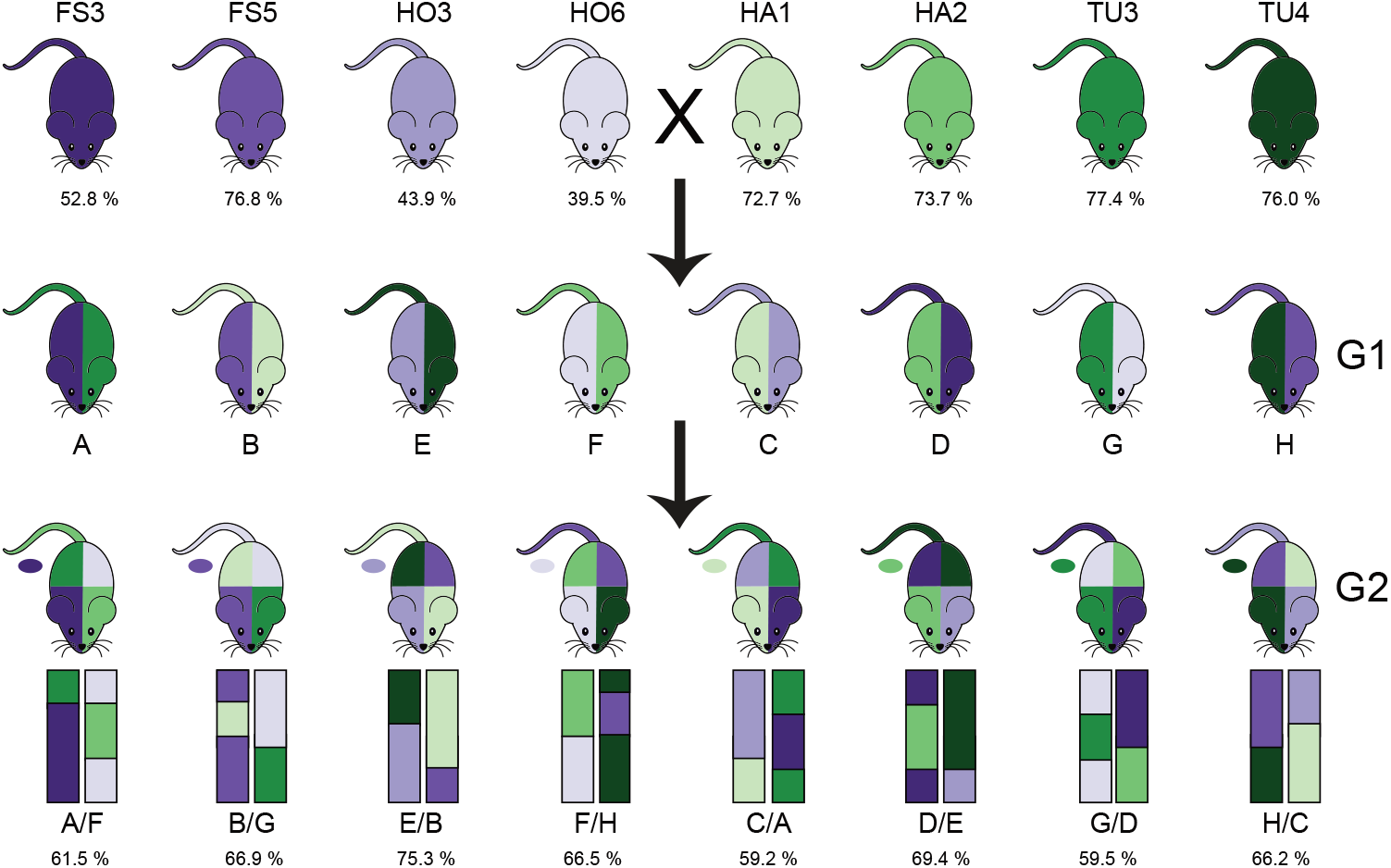
Overview of the intercross design. G0 mice are from eight partially inbred lines derived from mice wild-caught in four hybrid zone sites. Hybrid index - the percentage of *musculus* alleles - is reported as the mean for the G0 mice from each line (top), or mean of 40 G2s from each subcross (bottom). We performed eight G1 crosses with one line with hybrid index ~50% (purple shades) and one line with hybrid index >50% (green shades); color on the left side of mouse diagram indicates dam line and right side indicates sire line. Next, G1 mice were crossed in eight combinations such that each G2 mouse had one grandparent from each of the four breeding stocks, indicated by colors of mouse diagram, and representative chromosomes below. Tail color indicates Y chromosome strain, and oval indicates mitochondrial strain.

## Notes

https://github.com/sdoms/mapping_scripts

### Competing Interest Statement

The authors have declared no competing interest.

