## Supplementary figures for "Key features of the genetic architecture and evolution of host-microbe interactions revealed by high-resolution genetic mapping of the mucosa-associated gut microbiome in hybrid mice"

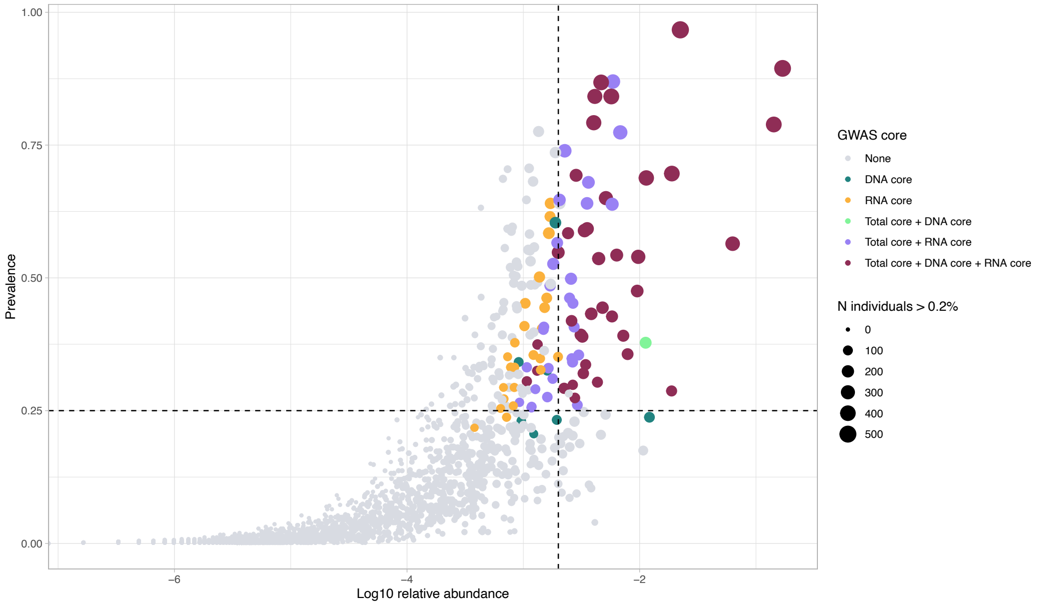


**Supplementary figure 1**: Selection of taxa for mGWAS analysis. A scatter plot showing the association of average relative abundance of taxa with their prevalence in the G2 mapping population. Taxa retained for analysis are colored according to the originating core. The size of each dot represents the number of individuals that have a median abundance higher than 0.2% of the taxon. The dashed lines represent the thresholds of the core (vertical: median abundance>0.2% and horizontal prevalence of 25 %.

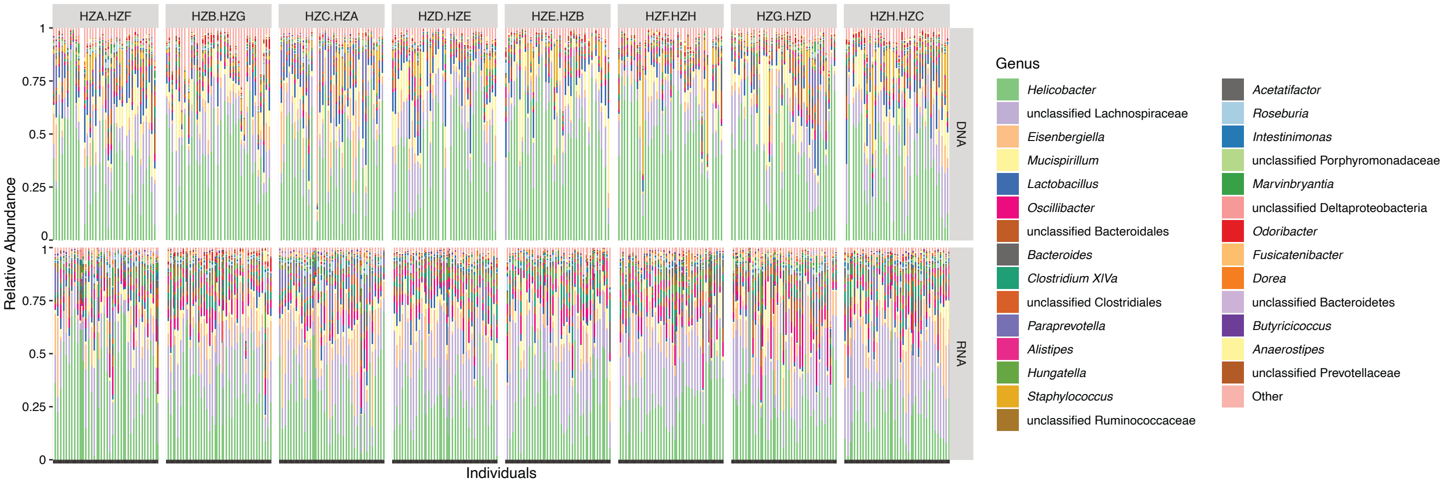


**Supplementary figure 2**: Relative abundances of core genera in G2 mapping population. Each vertical line represents one individual. Subcross (see supplementary figure 14) is indicated at the top.

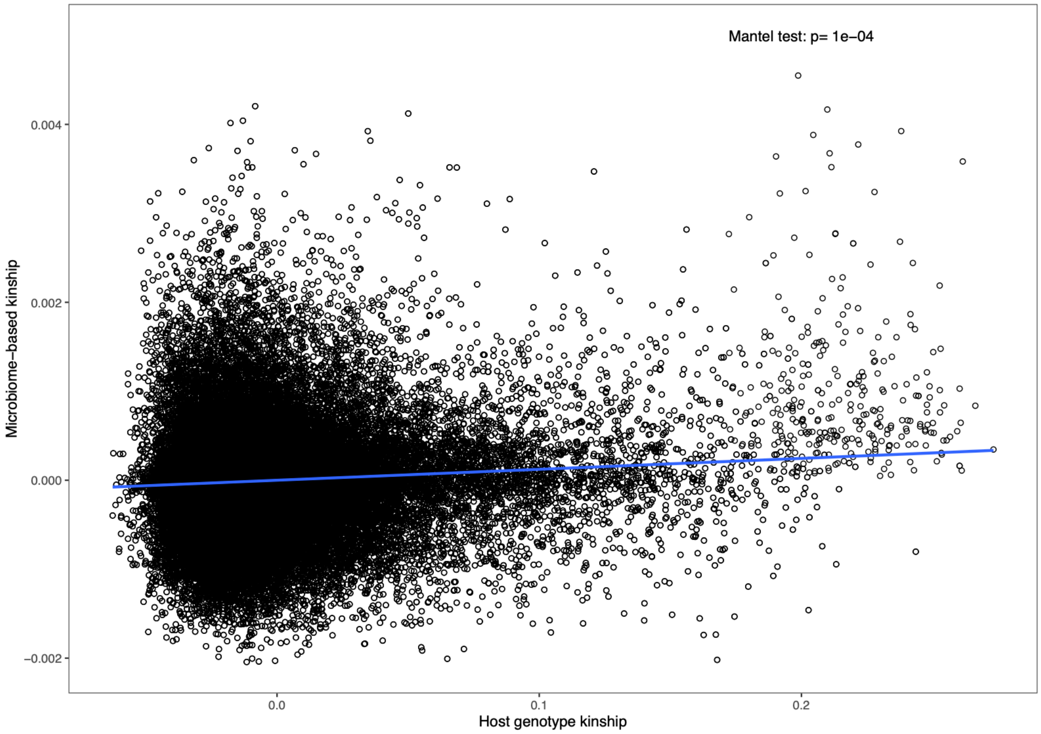


**Supplementary figure 3**: Host genetic relatedness calculated from SNP data (x-axis) correlated with microbial composition-based relatedness (y-axis) calculated from ASV abundances. The blue line represents a linear regression fit to the data.

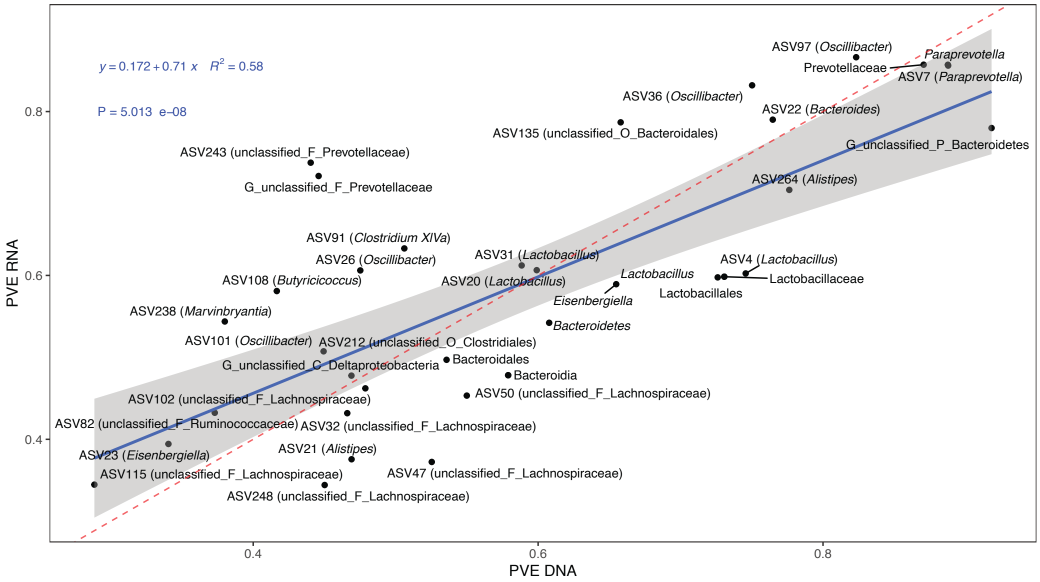


**Supplementary figure 4**: Correlation of SNP-based heritability estimates based on DNA (x-axis) or RNA (y-axis). The blue line represents a linear regression fit to the data. Red dashed line represents the identity line with a slope of 1.

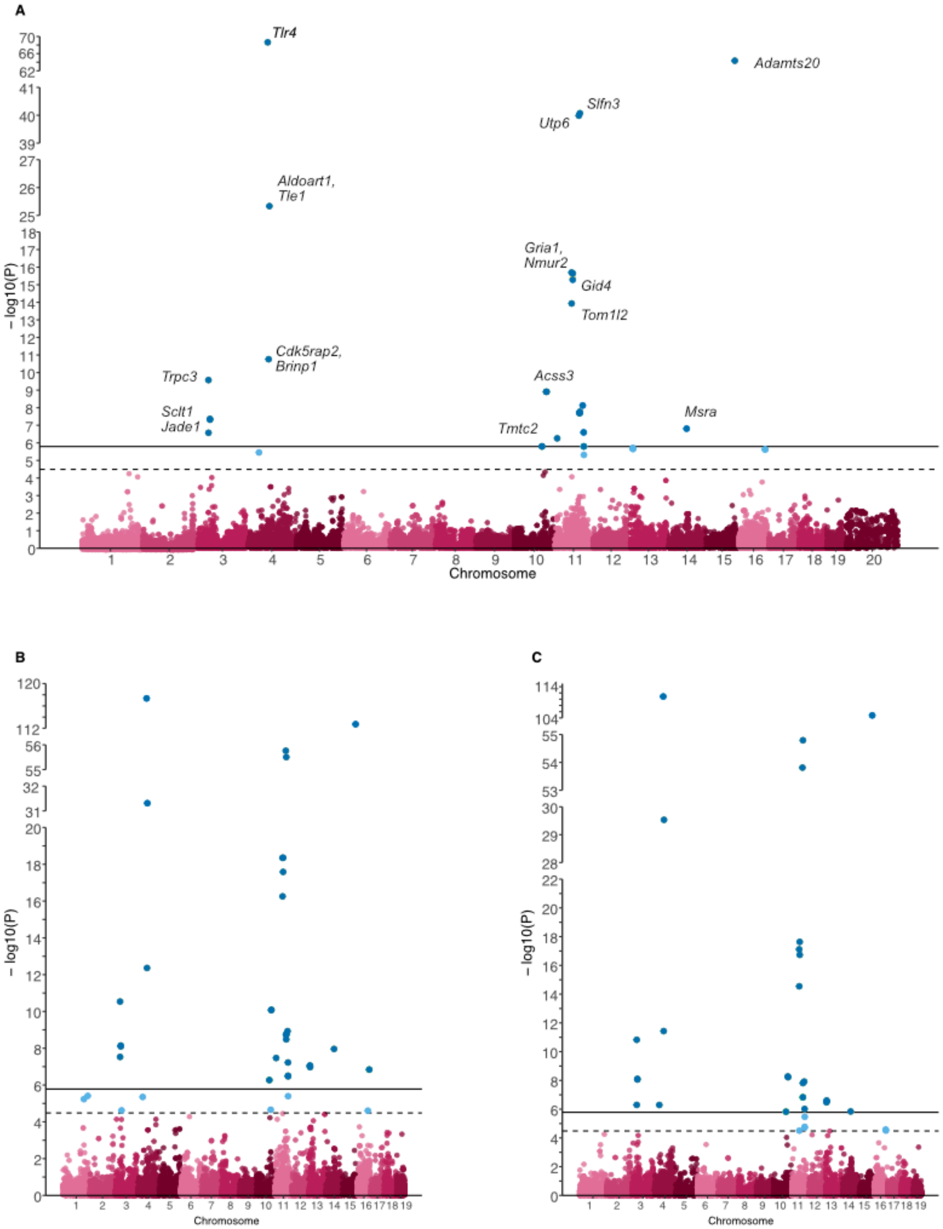


**Supplementary figure 5**: Manhattan plots for ASV184 (*Dorea)* of the complete model (A), the additive effect (B) or the dominance effect (C). SNPs passing the study-wide significance threshold (solid line) are shown in dark blue, while genome-wide significant SNPs (dashed line) are shown in light blue. In panel A, the closest gene to the SNP is shown for a subset of significant SNPs.

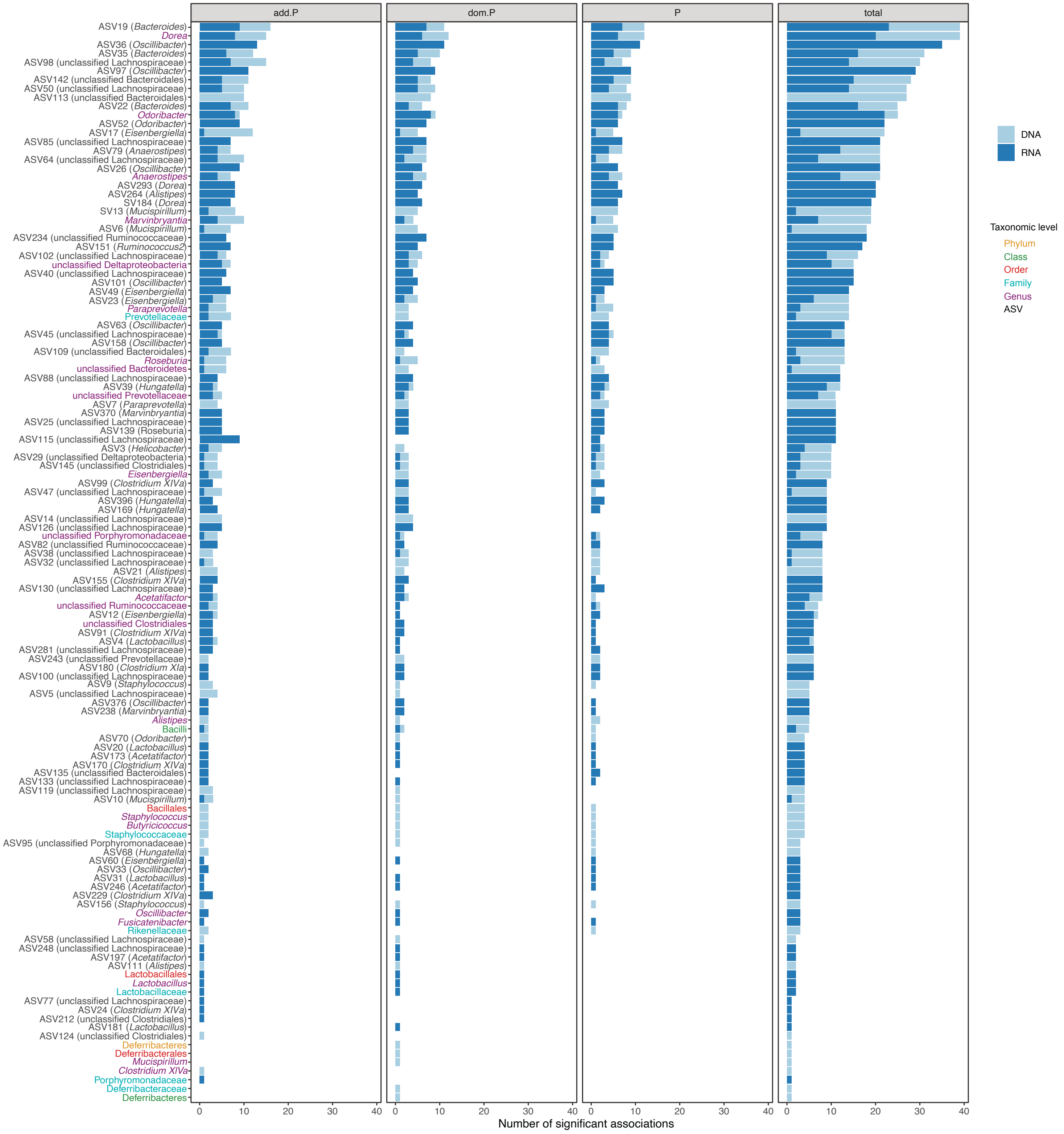


**Supplementary figure 6**: Number of significantly associated loci per bacterial taxon. Loci with significant additive effects (add.P), dominance effects (dom.P) or effects in full model (P) are indicated.

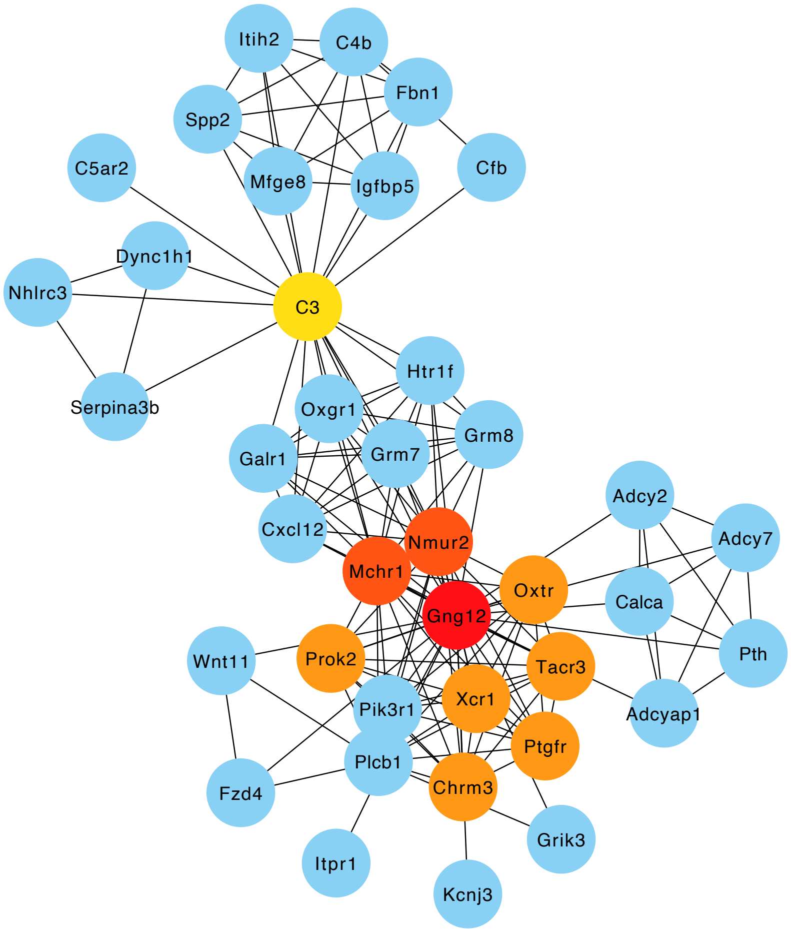


**Supplementary figure 7**: Top ten hub genes of the protein-protein interaction (PPI) network with the closest genes to the host SNPs significantly associated with bacterial abundances. The nodes are colored according to hub gene rank from 1 (red) to 10 (yellow). Blue nodes are the first neighbors.

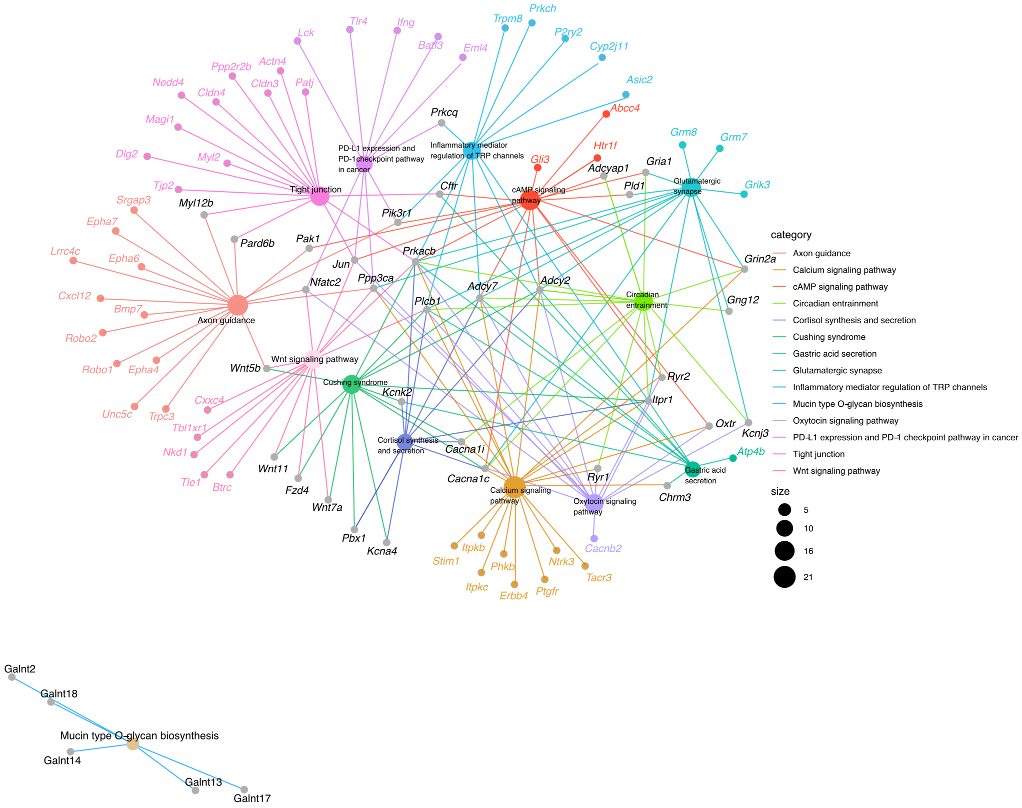


**Supplementary figure 8**: Genes belonging to over-represented KEGG pathways within the host genes closest to significant SNPs from association analysis.

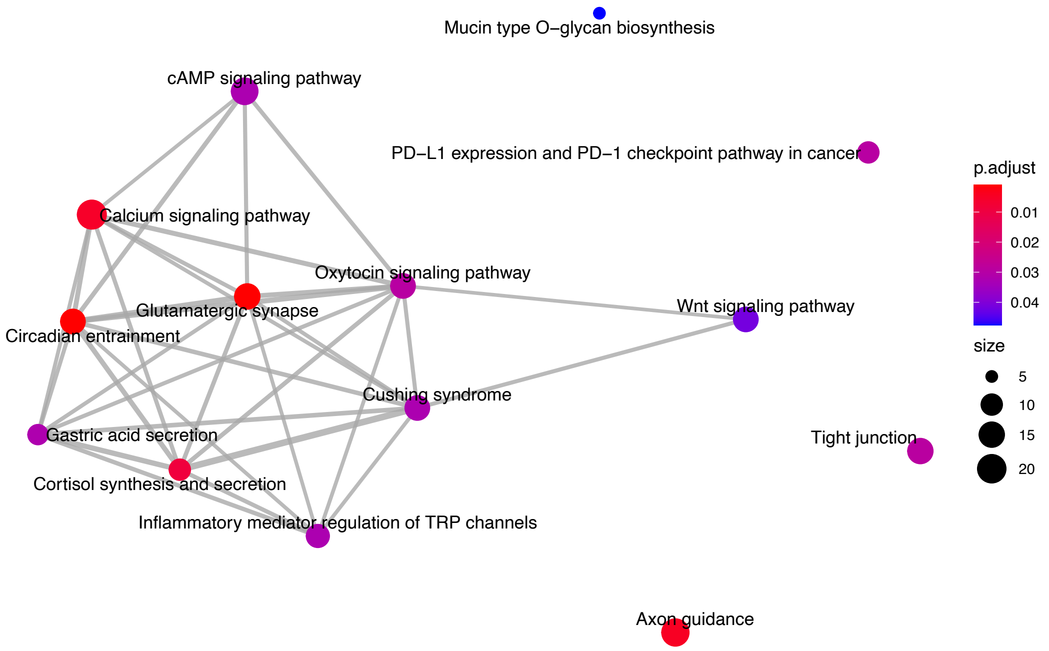


**Supplementary figure 9**: Enriched KEGG pathways among closest genes to significant SNPs.from association analysis. Node color indicates FDR-adjusted *P* value of enrichment and node size indicates number of candidate genes in pathway.

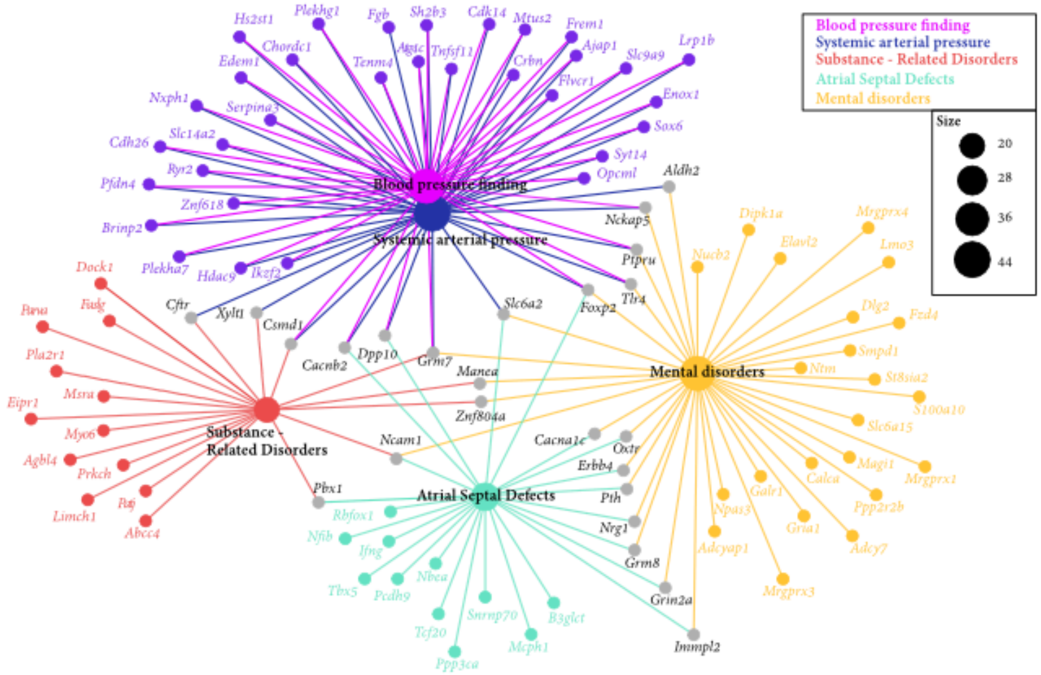


**Supplementary figure 10**: Enriched human diseases among genes closest to significant SNPs from association analysis.

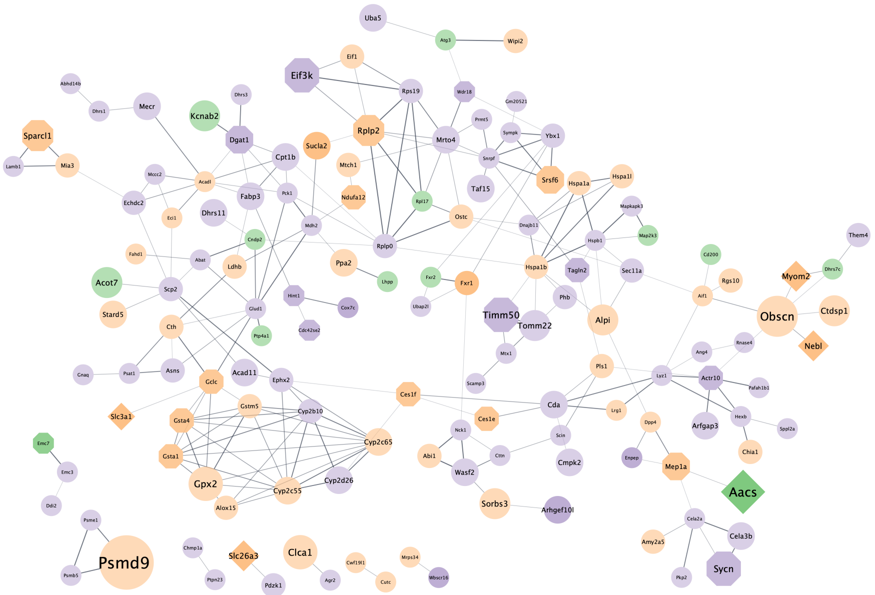


**Supplementary figure 11**: STRING (Szklarczyk et al., 2019) protein-protein interaction network of proteins that are differentially expressed in the intestine (small intestine and colon) of germ-free (GF) mice compared to conventionally raised mice, found in the present study. The color of the network nodes indicates whether the QTL hit was found using the DNA abundances (green), RNA abundances (purple) or was found in both (orange). The shape represents if the gene of the protein was the closest gene to the significant SNP (rectangle), if the gene was also found in QTLs of other studies (octagon), a combination of both (diamond), or only differentially expressed in GF mice vs. conventionally raised mice. The node size expresses the number of taxa where the gene was found in a QTL. The edges represent protein-protein interactions, where the line thickness indicates the strength of the data support from text mining, experiments, databases, co-expression, gene-fusion, and co-occurrence.

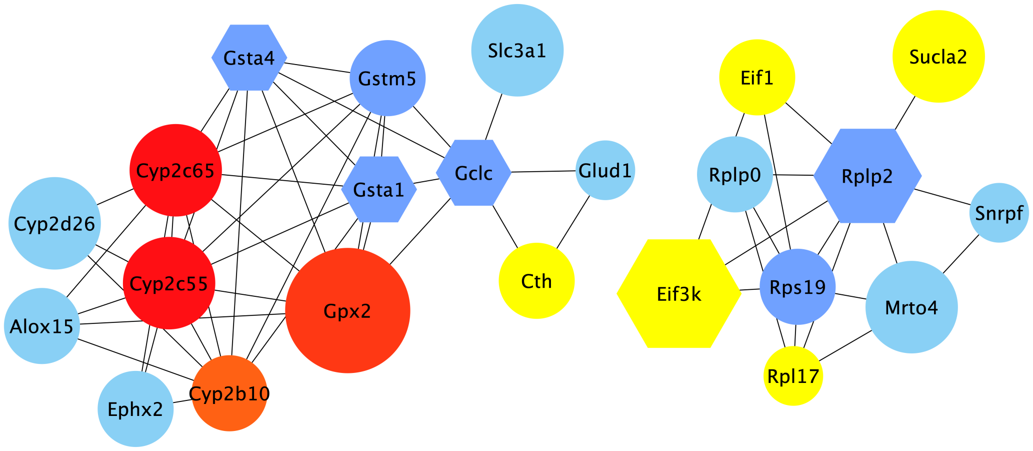


**Supplementary figure 12**: Visualization of the top hub genes calculated with the MCC algorithm and their first neighbors from the protein-protein interaction (PPI) network of genes found in intervals in present study that are also differentially expressed in germ-free versus conventionally raised mice. Edges represent the protein-protein associations. The red nodes represent genes with a high degree (= hub genes), and the yellow nodes with a low degree, while the blue nodes represent their first neighbors. All nodes shown are differentially expressed in GF mice. Hexagon shaped nodes are genes/proteins also found associated with gut microbiome abundances in other mouse QTL studies, and round nodes are 'only' differentially expressed in GF mice. The size of the node is an indication of the amount of taxa associated with the gene.

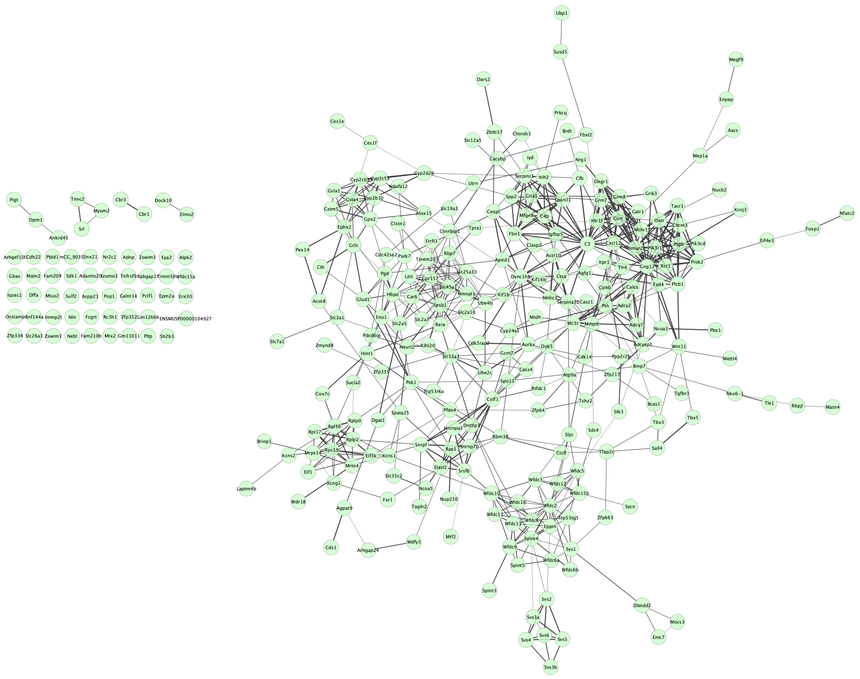

**Supplementary figure 13**: Original protein protein interaction (PPI) network of 304 candidate genes closest to SNPs significantly associated with bacterial abundances. Generated in STRING (Szklarczyk et al., 2019) and Cytoscape (Shannon et al., 2003).

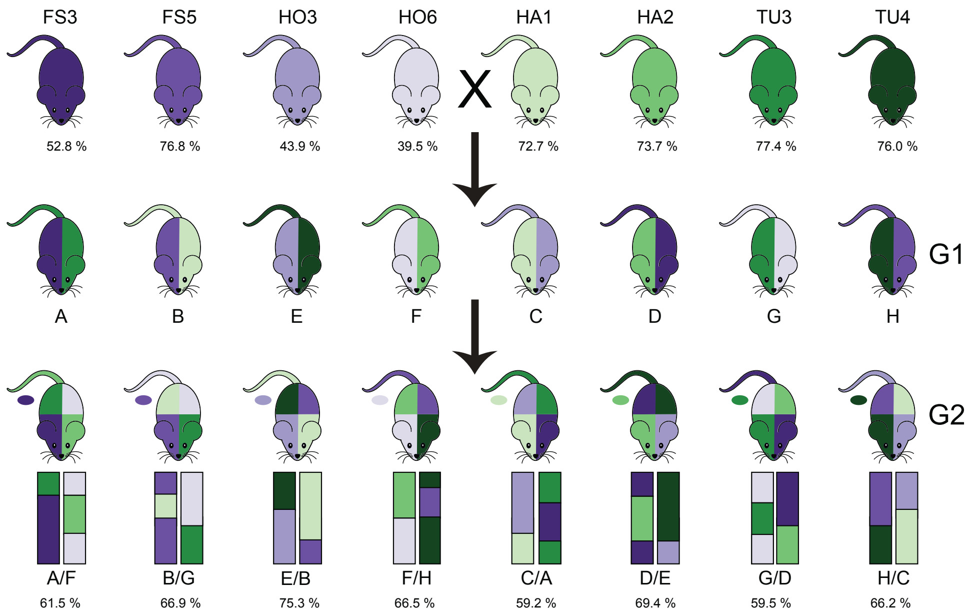


**Supplementary figure 14**: Overview of the intercross design. G0 mice are from eight partially inbred lines derived from mice wild-caught in four hybrid zone sites. Hybrid index - the percentage of *musculus* alleles - is reported as the mean for the G0 mice from each line (top), or mean of 40 G2s from each subcross (bottom). We performed eight G1 crosses with one line with hybrid index ~50% (purple shades) and one line with hybrid index >50% (green shades); color on the left side of mouse diagram indicates dam line and right side indicates sire line. Next, G1 mice were crossed in eight combinations such that each G2 mouse had one grandparent from each of the four breeding stocks, indicated by colors of mouse diagram, and representative chromosomes below. Tail color indicates Y chromosome strain, and oval indicates mitochondrial strain.
